# PolyA/polyQ-mediated conformational rewiring regulates DNA engagement and drives aggregation in the neuronal transcription factor Ascl1

**DOI:** 10.64898/2026.05.11.724289

**Authors:** Shrutarshi Mitra, Sarah F. Ruidiaz, Ásdís H.V. Laufeyjardóttir, Vasileios Voutsinos, Kateryna Nitsenko, Andrea Guala, Kenneth S. Zaret, Rasmus Hartmann-Petersen, Pétur O. Heidarsson

## Abstract

Ascl1 is a pioneer transcription factor that drives neuronal fate decisions, yet the structural basis of its activity remains elusive. Besides the basic helix–loop–helix (bHLH) domain which dimerizes with other transcription factors and binds DNA, Ascl1 contains long low-complexity intrinsically disordered regions (IDRs), including a polyA/polyQ tract of unknown function. Here we use single-molecule FRET to map the conformational ensemble of full-length Ascl1 across monomeric, heterodimeric, and DNA-bound states. Monomeric Ascl1 is largely disordered but displays multivalent coupling between the N-terminal IDR and the bHLH domain, with the polyA/polyQ tract opposing bHLH compaction. E12 binding folds the bHLH domain and remodels the N-IDR, increasing dynamics across the N-IDR while suppressing them locally within the polyA/polyQ tract. Deleting the tract weakens nonspecific DNA binding by the Ascl1/E12 heterodimer without disrupting heterodimerization or high-affinity E-box binding, abolishes aggregation in vitro, and increases Ascl1 abundance in HEK293T cells. Thus, the polyA/polyQ tract acts as a regulatory module that promotes nonspecific DNA engagement while imposing a cost in solubility and cellular abundance.

## INTRODUCTION

Achaete-scute homolog 1 (Ascl1) is a proneural transcription factor that initiates neurogenesis by driving the commitment of progenitor cells to a neuronal fate and activating neuronal differentiation programs. Ascl1 also functions as a pioneer transcription factor^1,2^, a class of transcription factors that possess distinct abilities to target DNA in condensed chromatin and initiate cell-fate changes^3,4^, and as such it can be used to drive fibroblasts to form induced neuronal cells (iN)^5^. However, a molecular understanding of the key players and mechanisms behind both neurogenesis and reprogramming is currently limited.

The 236-residue Ascl1 protein has a central basic helix-loop-helix (bHLH) domain (residues 118-185) flanked by intrinsically disordered regions (IDRs) (**Fig. 1A**). The bHLH domain has strong α-helical propensity but has been suggested to lack tertiary contacts in the free protein^6^ and to subsequently go through a disorder-to-order transition upon homo- or heterodimerization with other bHLH proteins. The long N-terminal region (residues 1-117, N-IDR) is a particularly interesting low-complexity IDR. Three quarters of the first 75 residues are composed of alanine (29%), glutamine (28%), proline (12%), or serine (7%), including a 13×alanine/12×glutamine (polyA/polyQ) tract of unknown function that forms at least a transient α-helix^6^ (**Fig. 1A**). As the hallmark of bHLH transcription factors, the basic region that binds to DNA and the HLH domain involved in dimerization are highly conserved. The N-IDR is less conserved, predominantly due to variations in and around the polyA/polyQ region length and composition, even in closely related species (**Supplementary Fig. S1**). Expansions of low complexity polyQ and polyA repeats, through codon duplication, have been associated with at least 18 different genetic diseases, and for Ascl1 a link has been established between polyQ length variation and Parkinsońs disease^7^. The shorter C-terminal IDR (C-IDR, residues 186-236) is predicted to contain a transcriptional activation domain^8^ and is rich in serine-proline pairs (**Fig. 1A**). Ascl1 preferentially forms heterodimers with the ubiquitously expressed bHLH factors called E-proteins, including TCF3 (previously E12/E47), TCF4, and TCF12^9^. Ascl1/E-protein heterodimers bind canonical Enhancer-box (E-box) consensus sites in DNA and activate transcription. While bHLH factors including Ascl1 have regions outside the bHLH domain essential for neurodevelopment^8,10^, how such non-DBD domains function is not known.

**Figure 1.**
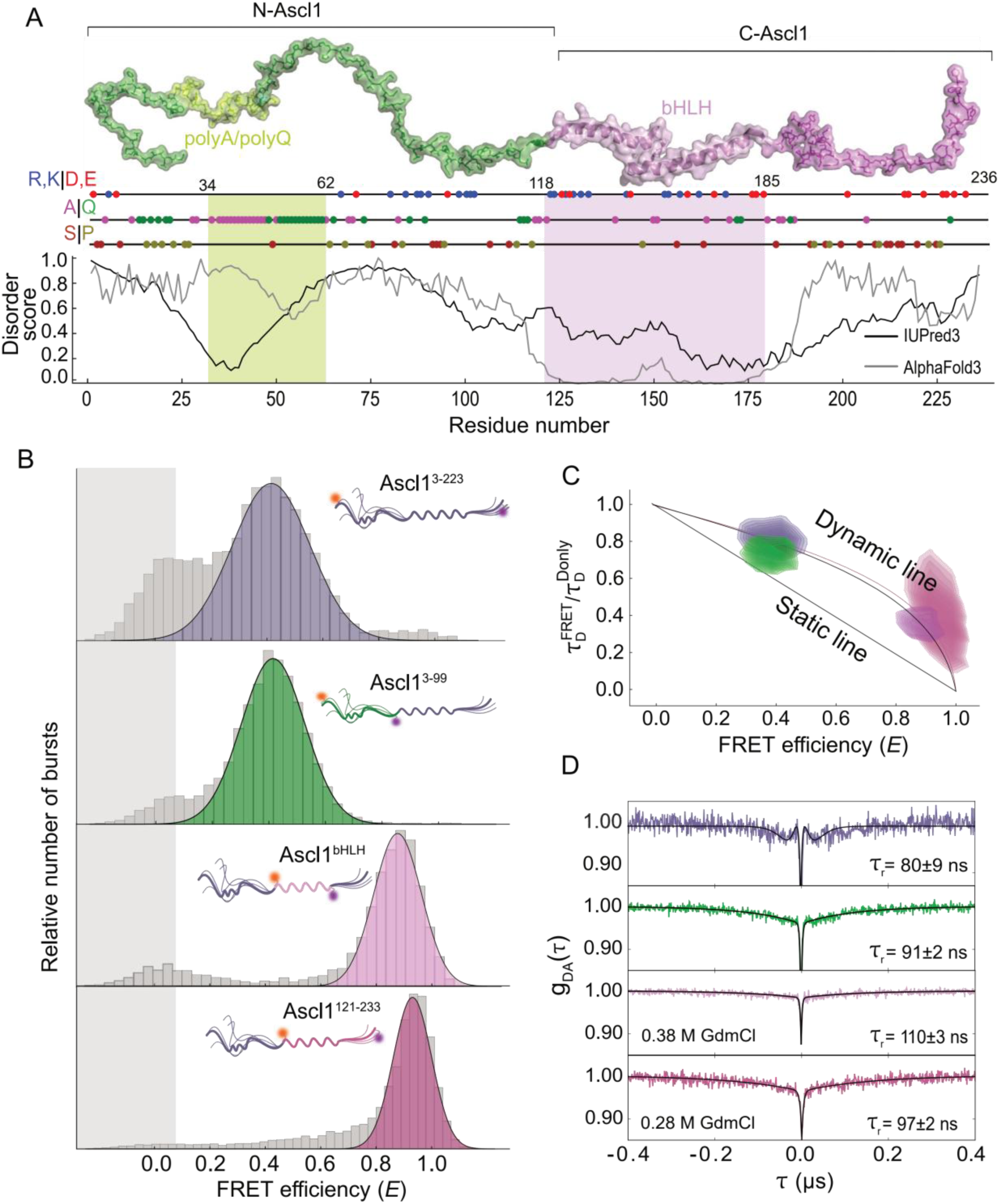
Monomeric Ascl1 is disordered and highly dynamic yet compact. **A)** Schematic of Ascl1 structure illustrating the major structural features: N-IDR (residues 1-117 including polyA/polyQ tract (green)), bHLH DNA binding domain (residues 118-185 (pink)), and C-IDR (residues 186-236 (magenta)). Below: Disorder prediction plot as a function of residue number, using AlphaFold3^24^ pLDDT scores (blue) and Disprot^25^ (black). Charged residues (Glu/Asp (red), Arg/Lys (blue)), as well as alanine/glutamine and serine/proline content are indicated. The same color scheme is applied in all subsequent figures. **B)** Single-molecule FRET efficiency histograms of Ascl1^3-233^ (blue), Ascl1^3-99^ (green), Ascl1^121-233^, and Ascl1^bHLH^ (pink). The peak at FRET efficiency ∼0 originates from residual donor-only molecules that remain after PIE filtering (**Supplementary Fig. S2**). **C)** Relative donor fluorescence lifetime analysis. Plots showing the fluorescence lifetimes of Cy3b 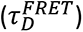 normalized by the intrinsic donor lifetime 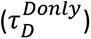 as a function ofFRET efficiency. The dynamic line is based on a SAW-ν polymer model^19^. **D)** Rapid reconfiguration dynamics probed with nsFCS. Solid lines are exponential fits and are used to determine the chain reconfiguration time, *τ*_r_. Due to the high FRET efficiency in the C-terminal constructs, the data shown includes small amounts of denaturant to aid visualizing the chain dynamics (**see Supplementary Fig. S5**).

Specifically, we lack an accurate description of full-length Ascl1 conformational ensemble, the potential interplay between ordered and disordered regions, and the nature of the conformational rearrangements that occur upon complex formation with E-proteins and DNA. The lack of structural and functional information on Ascl1 is not only due to the high degree of its inherent conformational disorder but also due to an extreme aggregation propensity of the full-length protein, impeding both its isolation and study^6^, and potentially limiting its application in neuronal reprogramming without a heterodimerization partner^11^. A quantitative description of full-length Ascl1 is therefore needed to understand how its IDRs influence dimerization, DNA binding, and self-association.

In this work, we conducted an in-depth characterization of the conformational ensembles of full-length Ascl1 in various molecular contexts using single-molecule spectroscopy in combination with Förster resonance energy transfer (smFRET)– a powerful technique to study disordered proteins^12–15^. The ultra-low detection limits of smFRET enabled us to use extremely low concentrations (picomolar) of Ascl1 in our experiments and avoid complications due to its high aggregation propensity. Using these techniques, we deciphered conformational dynamics across varying structural and functional states, including monomers, homo- and heterodimers, as well as complexes with different DNA constructs. Our results identify the polyA/polyQ tract as a conformational regulator that promotes nonspecific DNA binding and drives Ascl1 aggregation.

## RESULTS

### Ascl1 forms a compact and rapidly exchanging conformational ensemble

The high aggregation propensity of Ascl1 has thus far prevented structural characterization of the full-length protein^6^. Detection by smFRET requires mere picomolar concentrations of protein, allowing us to circumvent complications due to aggregation. We can thus probe Ascl1 dimensions and conformational dynamics within discrete polypeptide regions, including those that are disordered. We began by producing fluorescently labeled Ascl1 variants using the dyes Cy3b as donor and CF660R as acceptor (**Supplementary Tables 1, 2**). We produced full-length variants probing essentially the entire polypeptide chain or distinct subregions (**Fig. 1A**): End-to-end labeled (Ascl1^3-2^^33^, superscript indicates labeling position), N-terminal IDR labeled (Ascl1^3-99^), C-terminal half labelled (Ascl1^121-233^) and specifically labeled around bHLH domain (Ascl1^bHLH^: labeled on residues 121 and 181).

FRET efficiency between the donor and acceptor dye is highly dependent on the distance between them and can therefore be used as a sensitive molecular ruler in the 2-10 nm spatial range^16–18^. We measured FRET from thousands of individual molecules of each Ascl1 variant, and the resulting FRET efficiency histograms showed a single well-resolved population that was broader than expected from shot-noise-limited statistics, indicating conformational heterogeneity within the ensemble (**Fig. 1B**, **Supplementary Fig. S2, S3, Supplementary Table 2**). The high FRET efficiency of the C-terminal domain, including just the bHLH, is noteworthy, despite quite large sequence separation of the dyes (60-112 amino acids), and indicates that bHLH region populates a very compact conformation. By applying a self-avoiding walk (SAW) polymer model^19^, we determined the Flory scaling exponent, *ν*, which provides an estimate for compactness independent of sequence length. This analysis emphasizes how the two halves of Ascl1 have very different dimensions, with an extended N-terminal IDR (ν˃0.55) and a compact C-terminal with the bHLH domain (ν<0.45)^19,20^.

Fluorescence lifetime analysis can be used to detect distance fluctuations between donor and acceptor on a sub-millisecond timescale^21^. Such analysis for Ascl1 shows that the donor lifetimes are indeed positioned away from the static diagonal line (**Fig. 1C**, **Methods**), residing instead on the curved dynamic line, indicating a broad distance distribution characteristic of disordered proteins and IDRs. A slight shift towards the static line was observed with Ascl1^3-99^, indicating dampened dynamics in this region on the sub-millisecond timescale. Chain dynamics can be further quantified using fluorescence correlation spectroscopy (FCS) on the nanosecond timescale (nsFCS)^22,23^ which allows determining the reconfiguration time of the chain (*τ_r_*) (**Fig. 1D**, **Supplementary Fig. S4**, **S5, S6**). We observed rapid reconfiguration dynamics across the polypeptide chain (*τ*r=80-110 ns), representative of a highly disordered and dynamic protein, and in line with the fluorescence lifetime analysis. This analysis also revealed that the N-IDR contributed most of the observed dynamics, as it reconfigured on a similar timescale as the entire protein, whereas the bHLH domain displayed slightly slower and lower amplitude dynamics (**Fig. 1D**). Overall, our data shows that Ascl1 has a highly compact structure yet retains rapid structural dynamics across its sequence.

### The N-IDR and polyA/polyQ tract exert opposing effects on bHLH compaction

Despite the substantial structural disorder in Ascl1, direct interactions between domains may contribute to the overall compact structure. To probe interdomain interactions, we created deletion variants N-Ascl1^3-99^ (isolated N-terminal half) and C-Ascl1^bHLH^ (isolated C-terminal half) focusing specifically on the bHLH domain. Interestingly, the 〈*E*〉 values for both N-Ascl1^3-99^ and C-Ascl1^bHLH^ were substantially lower when compared with the full-length protein (〈*E*〉 ∼0.32 vs. 0.39 and 〈*E*〉 ∼0.77 vs. 0.88, respectively), indicating that intramolecular interactions between the two domains lead to increased compaction for both domains (**Fig. 2A**). Furthermore, sub-millisecond and nanosecond distance dynamics were now more pronounced in the N-IDR when the neighboring C-terminal was absent in N-Ascl1^3-99^, again indicating loss of dynamic restraints from intramolecular interactions in full-length Ascl1 (**Fig. 2B, C**).

**Fig. 2.**
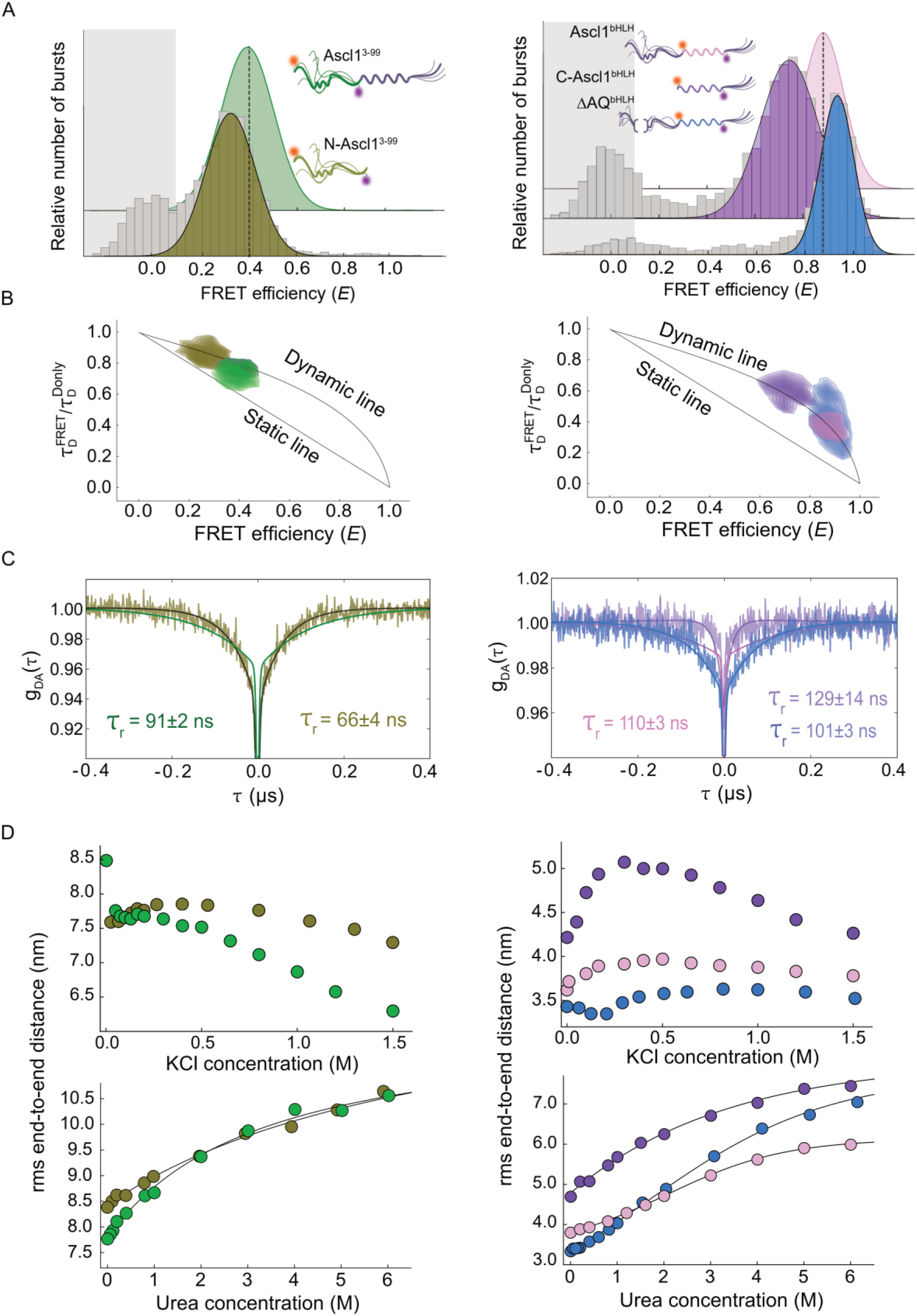
Deletion variants reveal interdomain effects on dimensions and stability of Ascl1. **A)** Single-molecule FRET efficiency histograms of N-Ascl1^3-99^ (lightgreen), C-Ascl1^bHLH^ (purple) and ΔAQ^bHLH^ (blue). **B)** Relative donor fluorescence lifetime analysis. Plots showing the fluorescence lifetimes of Cy3b 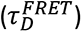 normalized by the intrinsic donor lifetime 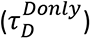 as a function of FRET efficiency. The dynamic line is based on a SAW-ν polymer model^19^. **C)** Rapid reconfiguration dynamics probed with nsFCS. Solid lines are exponential fits and are used to determine the chain reconfiguration time, *τ*_r_. The fit line for the full-length variants (**Fig. 1D**) is shown for comparison. **D)** RMS distance between dyes as a function of KCl or urea concentration for Ascl1 variants. The solid lines represent fits to a weak denaturant model or a cooperative unfolding transition.

To understand whether the polyA/polyQ tract was involved in interdomain interactions, we created a polyA/polyQ deletion variant with fluorophores still flanking the bHLH domain (ΔAQ^bHLH^). Remarkably, the polyA/polyQ deletion variant had an even higher FRET efficiency than the WT bHLH domain, with 〈*E*〉∼0.92 compared to ∼0.88, indicating increased compaction and thus seemingly contrasting effects compared to removing the entire N-IDR (**Fig. 2A**).

To investigate whether the interdomain interactions originated from charged residues, we studied the dimensions of Ascl1 variants while modulating salt concentrations (**Fig. 2D**). We measured FRET efficiencies and converted to root mean squared (*RMS*) distance^19^ as a function of salt concentration (**Supplementary Fig. S7A,B, S8)**. Interestingly, the two terminal halves within the full-length protein had contrasting responses to salt concentration, evident with dramatic compaction at low salts for the N-terminal IDR and a modest expansion for the C-terminal half. The salt response was then markedly changed for our deletion variants: N-Ascl1^3-99^ now expanded with salt while the bHLH domain expansion was exaggerated when the entire N-terminal IDR was removed. However, when only the polyA/polyQ tract was removed, the bHLH compacted with salt. This result indicates strong but complex coupling of the two domains through electrostatic interactions, with counteracting contributions within N-IDR subregions.

We then used chemical denaturants to assess the conformational stability of Ascl1 and subregions, and to detect persistent secondary or tertiary structure (**Fig. 2D**, **Supplementary Fig. S7A, C**). With increasing concentrations of urea, the peak in the FRET efficiency histogram of Ascl1^3-99^ shifted gradually towards lower FRET values but the denaturation profile for the bHLH domain appeared mildly sigmoidal, indicating a cooperative unfolding transition. With the entire N-terminal IDR removed, the isolated C-Ascl1^bHLH^ displayed smooth unfolding whereas only removing the polyA/polyQ tract (ΔAQ^bHLH^) led to more pronounced cooperative unfolding, in line with the effects on dimensions. These results imply that both electrostatic and hydrophobic interactions between the two domains are responsible for modulating dimensions. Adding up to 2 µM unlabeled N-Ascl1 to 100 pM fluorescently labeled C-Ascl1^bHLH^ showed little effect on the mean FRET efficiency, indicating a weak or tether-dependent interaction between the two domains (**Supplementary Fig. S9A, B)**. Overall, our results are consistent with dynamic long-range and multivalent intramolecular interactions involving the Ascl1 low-complexity IDR, including the polyA/polyQ tract, and the bHLH domain.

### The polyA/polyQ tract drives aggregation and limits cellular abundance

Given the pronounced effect of the polyA/polyQ tract on the Ascl1 conformational ensemble, we next asked whether this low-complexity segment also accounts for the extreme aggregation propensity of the full-length protein. Aggregation can be directly monitored in our single molecule experiments as aggregates typically contain multiple fluorophores and thus have clearly distinguishable fluorescence intensity compared to that originating from individual molecules. With only Ascl1^bHLH^ present at picomolar concentrations, a plot of photon counts over time with 1 s binning showed that the fluorescence intensity fluctuated homogeneously around a mean value but was stable over the course of the measurement, without noticeable intense bursts (**Fig. 3A**). When we added increasing concentrations of unlabeled Ascl1 to picomolar concentrations of fluorescently labeled Ascl1^bHLH^ and monitored the effects on fluorescence-time traces, we started to detect longer and more intense fluorescence bursts already at approximately 100 nM concentration (**Fig. 3A**). A clear concentration dependence in number of aggregates was observed in the fluorescence time traces with increasing concentration of unlabeled Ascl1. If these bursts were indeed due to large oligomers, we would expect them to display substantially increased diffusion times compared to small monomers. We therefore used FCS to probe the diffusion timescale of the components present in the mixture. Upon increasing the concentrations of unlabeled full length Ascl1, we observed a dramatic shift of the FCS curves towards longer times, consistent with Ascl1 self-association and increased molecular size for diffusing protein species (**Fig. 3B**).

**Figure 3.**
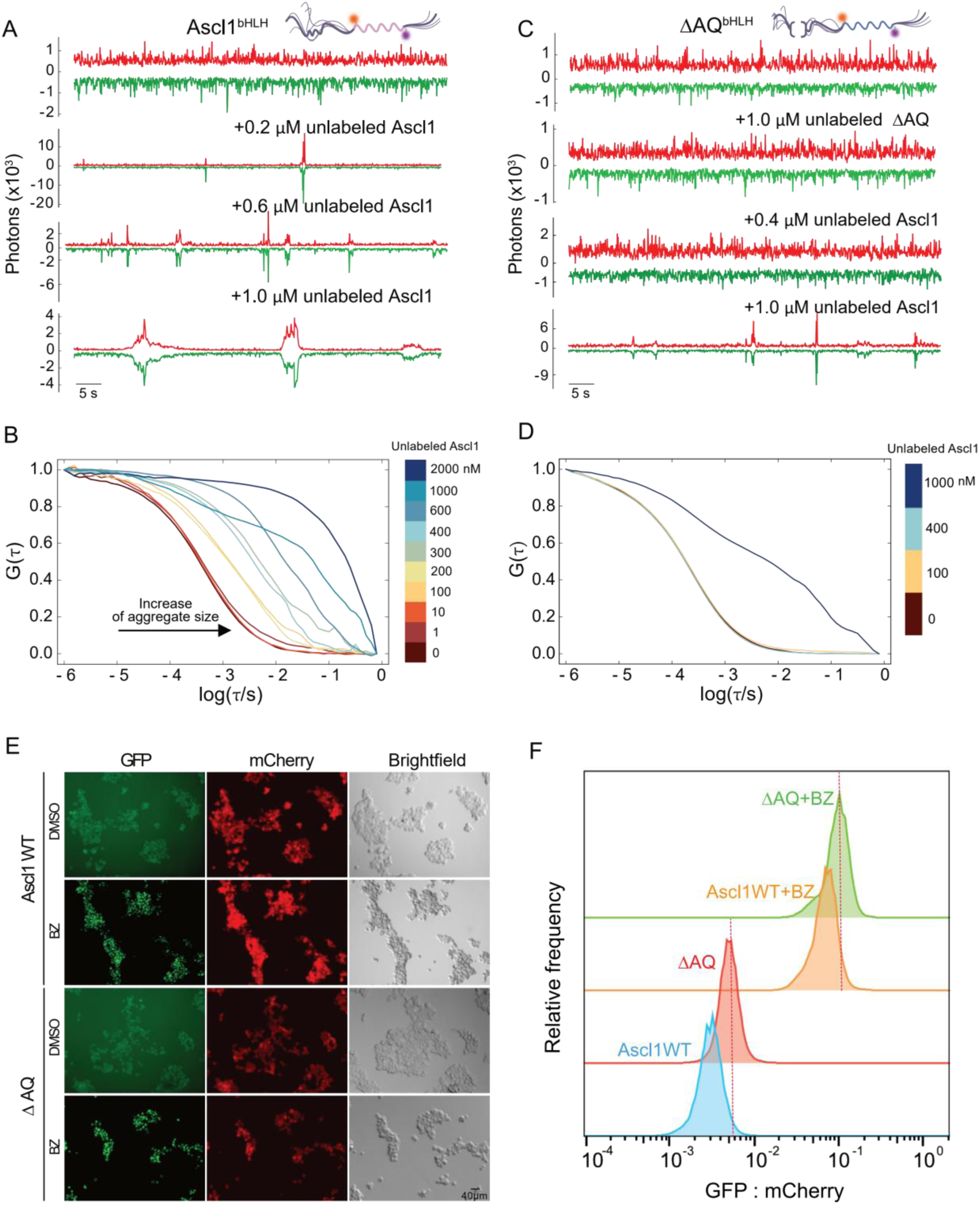
PolyA/polyQ tract drives extreme aggregation of Ascl1 and affects cellular abundance. **A)** Fluorescence vs time traces of 100 pM Ascl1^bHLH^ with 0-1 µM unlabeled WT Ascl1 with 1 s binning. **B)** FCS analysis of Ascl1^bHLH^ at varying concentrations of unlabeled WT Ascl1. **C)** Fluorescence vs time traces of 100 pM ΔAQ^bHLH^ with 1 µM unlabeled ΔAQ or 0.4 µM and 1 µM unlabelled WT Ascl1, with 1 s binning. **D)** FCS analysis of ΔAQ^bHLH^ at varying concentrations of unlabeled WT Ascl1. **E)** Fluorescence images of HEK293T cells expressing either Ascl1 WT or ΔAQ. Ascl1 was NLS-GFP tagged and mCherry was expressed from the same mRNA as NLS-GFP-Ascl1 for RNA levels normalization. **F)** Representative flow-cytometry distributions of GFP:mCherry ratios in cells expressing Ascl1 WT or ΔAQ, with or without bortezomib treatment (15 μM, 16 h). Data are shown from one of two independent experiments; *n* denotes the number of gated cells in the displayed experiment: WT, *n* = 12,958; ΔAQ, *n* = 12,819; WT + BZ, *n* = 12,692; ΔAQ + BZ, *n* = 12,073.

We then tested whether the polyA/polyQ tract contributes to the extreme aggregation propensity of Ascl1 by measuring the aggregation propensity for the polyA/polyQ deletion variant (ΔAQ^bHLH^). Remarkably, this variant was fully soluble even in the presence of micromolar amounts of its unlabeled counterpart, with no detectable aggregate bursts (**Fig. 3C**). We then investigated whether the ΔAQ^bHLH^ mutant would aggregate with high amounts of unlabeled WT Ascl1 that still contained the polyA/polyQ tract, to see if aggregation is due to polyA/polyQ tract interactions with other regions within the protein. We observed no change in the fluorescence intensity or protein diffusion time at nanomolar concentration of WT Ascl1 with only minimal aggregates observed at ∼1 µM (**Fig. 3D**). Measurements of solution turbidity at 350 nm were in excellent agreement with our FRET results (**Supplementary Fig. S10**) and showed that the polyA/polyQ deletion variant was highly soluble at >100 µM.

We then wanted to determine whether deleting the polyA/polyQ would influence its abundance in a cellular environment. For that we used the HEK293T cell line that allows expression of nuclear localization signal (NLS)-GFP-tagged Ascl1 with or without the polyA/polyQ tract from a bicistronic mRNA that also produces mCherry to normalize Ascl1 protein abundance to mRNA levels^26^. In our system, Ascl1 indeed exhibited nuclear localization and upon inhibition of proteasome by bortezomib we observed aggregation of both wildtype (WT) and ΔAQ (**Fig. 3E**, **Supplementary Fig. S11**). The aggregation of both WT and the ΔAQ is probably due to the overexpression of the protein in combination with the robust abundance increase as a result of bortezomib treatment (**Fig. 3E**). Quantification of the abundance levels (GFP:mCherry ratio) in cells by flow cytometry, showed that deletion of the polyA/polyQ tract resulted in substantially increased protein abundance (**Fig. 3F**). Together, these results show that the polyA/polyQ tract drives Ascl1 aggregation in vitro and is associated with lower Ascl1 abundance in HEK293T cells. The polyA/polyQ tract therefore imposes a substantial cost in solubility and cellular abundance.

### E12 binding folds the bHLH domain and remodels the N-IDR

We next asked how formation of the physiological Ascl1/E12 heterodimer reorganizes this conformationally coupled protein (**Fig. 4A**). It is well established that bHLH transcription factors heterodimerize with other bHLH proteins, and interaction network analysis has previously revealed the ubiquitously expressed E12 (also known as Tcf3) as an Ascl1 interacting partner^27^ that greatly enhances cellular reprogramming^11^. Structures of various bHLH dimers show a general architecture of two α-helical monomers that dimerize through a hydrophobic helix-loop-helix interface, forming a four-helix bundle^28,29^. We used AlphaFold3^24^ (AF3) to predict the structure of Ascl1, E12, and their complex (**Supplementary Fig. S12A,B**). The AF3 prediction is highly confident for the fold of the two bHLH domains but fails to confidently describe the IDRs, as expected.

**Figure 4.**
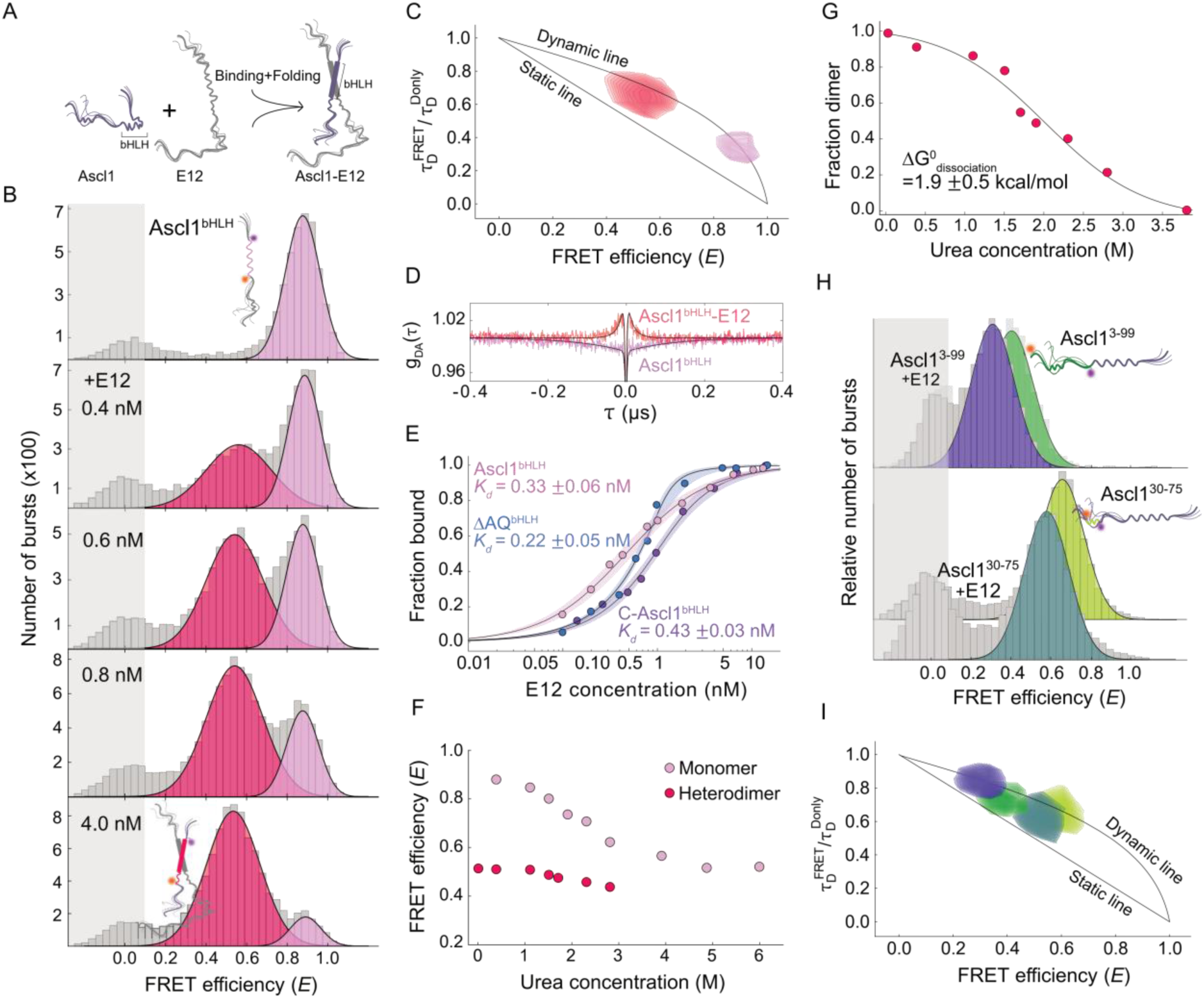
Conformational dynamics of Ascl1/E12 heterodimer. **A)** Schematic representation of monomer to heterodimer reaction. **B)** Single-molecule FRET efficiency histograms of Ascl1^bHLH^ with increasing concentrations of unlabeled E12. **C)** Relative donor fluorescence lifetime analysis and **D)** nsFCS of Ascl1^bHLH^ in complex with E12. **E)** Binding isotherms for the complex formation between Ascl1^bHLH^, ΔAQ^bHLH^, and C-Ascl1^bHLH^ with E12**. F)** FRET efficiency of heterodimeric complex of Ascl1^bHLH^ with E12 as a function of urea concentration. **G)** Fraction of heterodimer as a function of urea concentration. The solid line shows a fit to a two-state unfolding model (Methods). **H)** Single-molecule FRET efficiency histograms of Ascl1^3-99^ (upper) and Ascl1^polyA/polyQ^ (lower), free and in complex with 15 nM unlabeled E12. **I)** Relative donor fluorescence lifetime analysis of Ascl1^3-99^ and Ascl1^polyA/polyQ^, free and in complex with E12.

**Figure 5.**
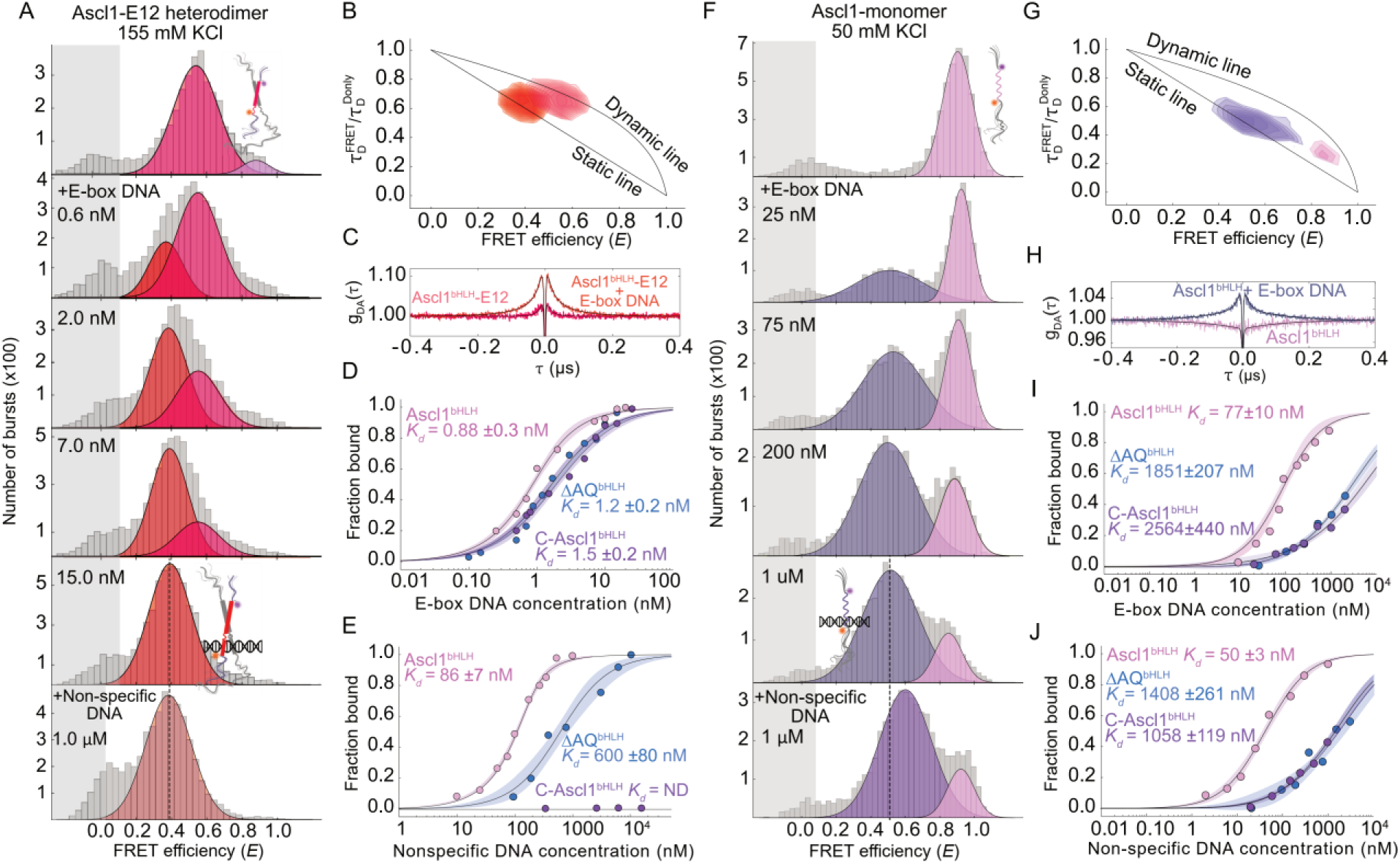
DNA binding of Ascl1 is modulated by the low-complexity IDR. **A)** Single-molecule FRET efficiency histograms of Ascl1^bHLH^/E12 complex at different DNA^E-box^ concentration and bound to DNA^nonspecific^ (bottom). **B, C)** Relative donor fluorescence lifetime analysis and nsFCS of Ascl1^bHLH^/E12/DNA complex. **D, E)** Binding isotherms for Ascl1^bHLH^/E12 (magenta), C-Ascl1^bHLH^/E12 (purple), and ΔAQ^bHLH^ (blue) as a function of DNA^E-box^ and DNA^nonspecific^ concentration. The solid lines represent fits to a 1:1 binding model. **F)** Single-molecule FRET efficiency histograms of monomeric Ascl1^bHLH^ binding to DNA^E-box^ and bound to DNA^nonspecific^ (bottom). **G, H)** Relative donor fluorescence lifetime analysis and nsFCS of monomeric Ascl1/DNA complex. **I, J)** Binding isotherms of monomeric Ascl1^bHLH^ (magenta), C-Ascl1^bHLH^ (purple), and ΔAQ^bHLH^ (blue) with DNA^E-box^ and DNA^nonspecific^. The solid lines represent fits to a 1:1 binding model.

**Fig. 6.**
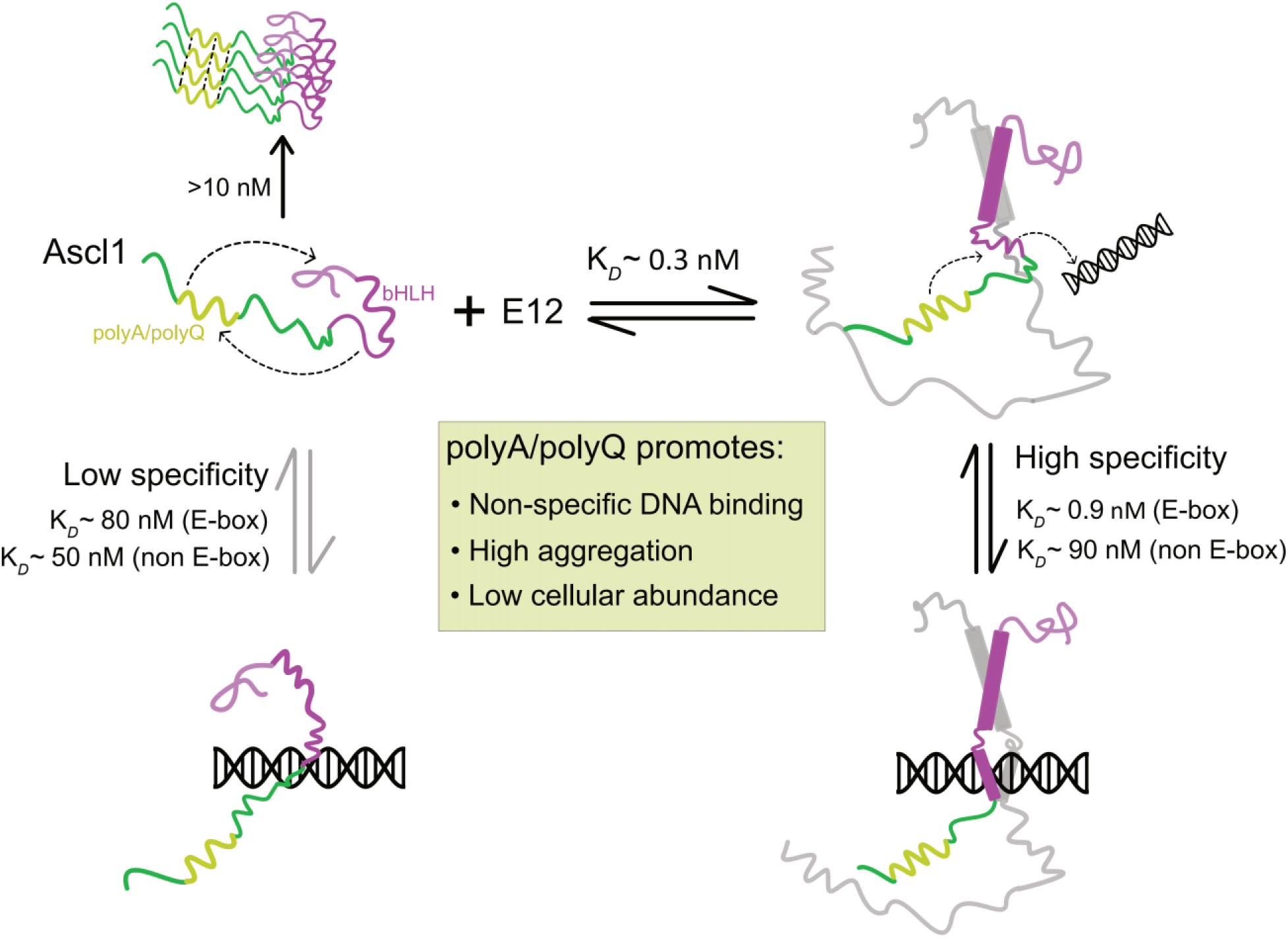
Model for polyA/polyQ-dependent regulation of Ascl1. The global dimensions of Ascl1 are based on our FRET mapping using six different dye pair locations that report on its structure in different contexts. Monomeric Ascl1 has competing pathways involving heterodimerization, nonspecific and specific DNA binding, and aggregation. The low-complexity polyA/polyQ tract plays a substantial role in mediating many of Ascl1’s molecular properties.

We then used our fluorescently labeled variants to probe structural changes in Ascl1 upon complex formation with E12. Ascl1 displayed a global expansion upon dimerization with E12 (**Fig. 4A,B**, **Supplementary Fig. S12C,D,E**). Complex formation can be confirmed by FCS analysis as the increased molecular size leads to almost 50% slower translational diffusion of the complex through the confocal volume (**Supplementary Fig. S12F**). Since the Ascl1 bHLH domain is known to engage with the bHLH domain of E12, we titrated Ascl1^bHLH^ with increasing concentrations of unlabeled E12. A large change in FRET was detected in presence of E12 from 0.88 to 0.47, indicating that the bHLH domain adopts a substantially more extended conformation in complex with E12, with ∼6.1 nm between the dyes, very close to the AF3 prediction (**Fig. 4B**, **Supplementary Table 3**). Furthermore, in the complex, the bHLH domain had substantially restricted sub-millisecond dynamics when probed with fluorescence lifetimes and nsFCS, as expected due to its folding into a well-defined structure (**Fig. 4C, D**). The FRET histograms of Ascl1 with varying concentrations of E12 directly report on the equilibrium populations and thus the fraction of bound protein. We therefore used the areas of the resulting populations in our FRET histograms to construct a binding isotherm for the complex formation with E12, allowing us to determine the binding affinity to E12, or *K*_d_ = 0.33 ± 0.06 nM– a very tight interaction (**Fig. 4E**, **Supplementary Table 3, 4**). Complex formation is dependent predominantly on hydrophobic interactions, since performing the same set of experiments at lower and higher salt concentration (100 mM and 300 mM KCl) yielded very similar dissociation constants (**Supplementary Fig. S12G**). We further investigated the stability of the dimer structure in the presence of urea and chose conditions such that Ascl1 is complexed in a stable dimer configuration (in the presence of 15 nM unlabeled E12 protein). The dimer complex showed some resistance to denaturation which can be visualized as an apparently cooperative transition representing dimer dissociation, with a free energy of unfolding *ΔG^0^*= 1.9 ± 0.53 kcal/mol (**Fig. 4F, G**).

Given that we observed long-range intramolecular interactions for monomeric Ascl1 (**Fig. 2**), we hypothesized that complex formation with E12 would affect the structure and dynamics of the N-terminal IDR of Ascl1. Ascl1^3-99^ in monomeric form had a rather broad FRET population with a mean transfer efficiency of ∼0.39 (**Fig. 1B**) and indeed, upon addition of unlabeled E12, a new population appeared with lower mean transfer efficiency of ∼0.32 (**Fig. 4H**). Fluorescence lifetime analysis showed that the addition of E12 shifted lifetimes towards the dynamic line compared to the free protein, indicating release of dynamic restraints from interdomain interactions (**Fig. 4I**), similar to removing the C-terminal half (**Fig. 2B**). We detected no independent interaction between the isolated N-IDR and E12 under these conditions (**Supplementary Fig. S12H**), and removing the N-terminal IDR had no effect on the binding affinity of C-Ascl1^bHLH^ to E12 (**Fig. 4E**). To probe specifically the polyA/polyQ tract, we produced a new Ascl1 variant, Ascl1^polyA/polyQ^, where the dyes were placed closely flanking the tract (**Fig. 4H**). Any changes in FRET efficiency upon complex formation would therefore almost exclusively reflect structural changes in the polyA/polyQ tract. We observed a distinct FRET change in this region from 0.6 to 0.53 upon binding E12 (**Fig. 4H**), indicating that the tract indeed changed conformation in the complex. Donor lifetime analysis and nsFCS revealed that rapid chain dynamics in the tract decreased in amplitude and reconfiguration time upon binding E12 (**Supplementary Fig. S5**), contrary to that observed for the entire N-terminal IDR. These results, combined with the differences in FRET and chemical stability between full-length Ascl1 and the N-terminal IDR missing the bHLH region (**Fig. 2**), support a model in which the polyA/polyQ tract contacts or otherwise conformationally couples to the bHLH region. To summarize, heterodimerization of Ascl1 with E12 both folds the bHLH domain and modifies the structure and dynamics of the N-terminal IDR.

### The polyA/polyQ tract promotes nonspecific DNA binding by Ascl1/E12

The bHLH transcription factor dimers preferentially bind promotor regions that contain E-Box sequences with the CANNTG hexamer as a consensus motif, where N can be any nucleotide but preferably C or G. Upon addition of 15 bp double-stranded DNA with an E-box sequence (DNA^E-box^, **Supplementary Table 4**), complex formation could be observed directly in our FRET efficiency histograms of the Ascl1^bHLH^/E12 complex. Complex formation with DNA led to an emergence of a third population with a 〈*E*〉 value of ∼0.38 (**Fig. 5A**, **Supplementary Table 3**), substantially lower than the 〈*E*〉 value of the free Ascl1^bHLH^/E12 complex (〈*E*〉 ∼0.47). DNA binding of bHLH proteins has been shown to stabilize helical structure at the basic region of the bHLH domain by increasing propensity for secondary structure^30^. Since in our experiment the dyes on Ascl1 are positioned N- and C-terminally of the bHLH region, the added formation of helix in the basic region increased further the donor-acceptor distance, causing the additional reduction in FRET efficiency, and suppressed even further distance fluctuations rendering them essentially static (**Fig. 5B, C**). SmFRET thus allows us to clearly resolve monomers, heterodimers, and their ternary DNA complexes, at equilibrium. Like before, we can use the areas of the populations with increasing amount of DNA to construct a binding isotherm and determine the DNA binding affinity to *K*_d_ = 0.88 ± 0.3 nM (**Fig. 5D**, **Supplementary Table 4**). E12 can in principle homodimerize and bind DNA itself but since its dimerization affinity is much weaker, we can exclude E12 DNA binding contribution from our analysis^31^.

The conformations of Ascl1 bHLH domain in the heterodimer with E12 seem to be identical when bound to DNA^E-box^ and non-specific DNA (DNA^nonspecific^, **Fig**. **5A**) but the binding affinity for non-specific DNA was ∼100-fold lower (**Fig. 5D, E**, **Supplementary Table 4**). To assess possible contributions from the N-IDR to DNA binding, we repeated DNA binding experiments using both C-Ascl1^bHLH^ (lacking the entire N-IDR) and ΔAQ^bHLH^. The binding affinity to E-box DNA was essentially unchanged for both variants compared to the full-length protein (**Fig. 5D**, **Supplementary Table 4**). Remarkably, non-specific DNA binding of ΔAQ^bHLH^ was severely compromised, with a dissociation constant close to the micromolar range ((*K*_d_= 600 ± 80 nM), and fully undetectable for C-Ascl1^bHLH^ even at concentrations of DNA up to 30 µM (**Fig. 5E**). Thus, the Ascl1/E12 heterodimer showed approximately 100-fold selectivity for E-box over nonspecific DNA. Removing the N-IDR left E-box affinity largely unchanged but abolished detectable nonspecific binding, whereas deletion of the polyA/polyQ tract weakened nonspecific binding approximately sevenfold. The absence of a detectable FRET response from the isolated N-IDR suggests that its contribution is mediated at least partly through conformational modulation of the bHLH domain but weak or heterogeneous N-IDR–DNA contacts cannot be excluded (**Supplementary Fig. S13**).

While most bHLH proteins require dimerization to bind DNA effectively, the cellular environment is likely to consist of a monomer-dimer equilibrium and some mechanistic studies even support a monomer-first binding pathway in certain contexts where monomers dimerize while bound to DNA^9,32^. We therefore studied whether an Ascl1-DNA^E-box^ complex could be observed under conditions where we only populate monomeric Ascl1. We could not observe any signs of complex formation upon addition of DNA at 165 mM salt but by reducing the KCl concentration to 50 mM (60 mM total ionic strength), a new and substantially broadened population emerged with a mean FRET efficiency of 0.52, higher than for the ternary complex with the heterodimer (0.37) and suppressed distance dynamics (**Fig. 5F, G, H**, **Supplementary Fig. S13**). To check if this DNA binding is a classical non-specific protein-DNA interaction we tried the same DNA binding assay with a non-specific DNA of same length (15bp) and a population at a slightly higher FRET efficiency emerged (**Fig. 5F**, **bottom**). We then performed careful titrations by adding varying amounts of unlabeled DNA and fitted the corresponding binding isotherms to determine the *K*_d_ (**Fig. 5I, J**). The binding affinity was similar for specific and non-specific DNA, 77 ± 10 nM and 50 ± 3 nM, respectively, or about 50-fold lower affinity than for the heterodimer. The low DNA binding affinity of the monomer and low degree of homodimerization indicates that DNA binding mostly occurs via the heterodimer pathway, as reported previously^33^. Large FRET changes were also observed in the N-IDR upon binding DNA (**Supplementary Fig. S13**) and we thus repeated binding affinity experiments with the construct lacking the N-terminal IDR or the polyA/polyQ tract. Under these low-salt conditions, monomeric Ascl1 bound E-box and nonspecific DNA with similar affinities, indicating little sequence discrimination. Removing either the entire N-IDR or only the polyA/polyQ tract weakened binding to both DNA sequences by more than an order of magnitude, demonstrating that the IDR also promotes DNA binding by monomeric Ascl1.

## DISCUSSION

In this study, we presented a conformational dissection of the pioneer transcription factor Ascl1, revealing a complex interplay linking its low-complexity IDRs, dimerization behavior, DNA binding properties, aggregation propensity, and cellular abundance. Interactions between IDRs and DNA binding domains have been observed before^34,35^, with varying effects on structure and function, but the strong contributions from the low-complexity polyA/polyQ tract that we observed for Ascl1 were unexpected. Even though monomeric Ascl1 is intrinsically disordered and highly dynamic, it exhibits a compact global architecture driven by the N-IDR engaging in multivalent but weak interactions with the bHLH domain. Removal of the entire N-IDR releases these contacts, resulting in a substantially more extended and salt-sensitive bHLH domain, consistent with electrostatically driven expansion of an unconstrained polypeptide. In contrast, selective deletion of the polyA/polyQ tract produced the opposite effect, compacting the bHLH, increasing its stability, and reversing its salt response, implying that the polyA/polyQ segment counterbalances other regions of the N-IDR that promote compaction. Together, these findings support a model in which the polyA/polyQ tract contributes to long-range conformational coupling within monomeric Ascl1, promotes nonspecific DNA engagement in its functional complexes, and simultaneously renders the protein strongly aggregation-prone. E12 binding reorganizes this ensemble by folding the bHLH domain and remodelling the N-IDR, and increasing its rapid nanosecond-microsecond dynamics but suppressing polyA/polyQ tract dynamics on the same timescale (**Fig. 6**).

Our FRET results revealed a new function for low complexity polyA/polyQ regions in facilitating DNA interactions. With the polyA/polyQ tract deleted, both monomeric Ascl1 and its heterodimeric complex with E12 underwent a similar DNA-induced folding of the bHLH domain, even though it displayed reduced nonspecific DNA affinity by almost an order of magnitude. That implies that the polyA/polyQ tract most likely reshapes the unbound ensemble and/or the binding pathway, rather than altering the overall geometry of the bound state. The selective effects on E-box DNA and nonspecific DNA are intriguing and suggest that with E-box DNA, specific contacts dominate the binding affinity. With nonspecific DNA, the N-IDR may increase the capture radius, promote transient interactions that increase productive encounters, or interact directly with DNA with weak heterogeneous contacts. Evolving modules that enable nonspecific binding may be particularly important for a pioneer transcription factor, as it enables it to effectively scan chromatin and potentially recognize nucleosomal structures^36,37^.

Finally, our results revealed the polyA/polyQ tract to be the main molecular driver that renders Ascl1 dramatically aggregation prone. Glutamine-rich domains are often found in transcription factors and have been suggested to be important for molecular recognition and to help with transcriptional activation^38^. Repeat expansions in glutamine- or alanine-rich tracts are well-known to drive aggregation in neurodegenerative disorders such as Huntington’s and Spinocerebellar ataxias^39,40^, and notably, Ascl1 polyQ length has been linked with risk to developing Parkinson’s disease^7^. The structure of polyQ sequences has been investigated in multiple biophysical studies^41,42^ and shown to display very different structural behaviour largely depending on the preceding amino acids sequences^43^. The polyA/polyQ tract of Ascl1 is highly variable in length across mammals, suggesting a tunable module for balancing function and stability (**Supplementary Fig. S1**).

The N-IDR of Ascl1 has been shown to mediate repression of mesendoderm genes^44^ and is likely to have other specific functions, yet the polyA/polyQ region has been shown to be redundant for neuronal differentiation^8^. That result raises an important evolutionary question—why maintain a highly aggregation-prone segment that is not strictly required for Ascl1 function? The answer probably involves the multiple roles of IDRs^45^, including enabling non-specific binding as we showed in this work. Another role for the polyA/polyQ tract may be regulatory, acting as a degron or aggregation “switch” that limits protein lifetime or local concentration in cells^46^. In a recent high-throughput study^26^, and recapitulated in this work, Ascl1 was shown to have very low abundance in human cells, implying rapid degradation. However, based on the peptide abundance predictor (PAP)^47^, the peptide consisting of the polyA/polyQ tract had only a weak degron score whereas the bHLH domain had a high score. A possible reason could be that the deletion of the polyA/polyQ tract removes its interactions with the bHLH domain, leading to its increased compactness which renders it less accessible to E3 ligases that target Ascl1. Why deletion of the tract increases cellular abundance remains unresolved; altered solubility, degradation, or both may contribute. Together, our results identify the polyA/polyQ tract as a sequence module that promotes nonspecific DNA engagement while increasing aggregation propensity and limiting cellular abundance.

## MATERIALS AND METHODS

### Protein expression and purification

Full length wild type (WT) Ascl1, as well as an N-terminal fragment (residues 1-117) and C-terminal fragment (residues 118-236), were designed according to Baronti et al.^6^ and cloned into a pET24b vector containing a 6×His SUMO-tag. The vector contains codes for a hexahistidine small ubiquitin-like modifier (His_6_-SUMO) tag added to the N-terminal of all constructs. Various cysteine mutants (after substituting the two native Cys30 and Cys99 with serines) were made using the QuikChange Lightning kit from Agilent using primers from Integrated DNA Technologies (IDT). All mutations and truncated constructs were verified using sequencing. Constructs were expressed in Lemo21(DE3) cells (New England BioLabs) cultured in LB-broth medium. Expression was induced at OD_600_ 0.5–0.7 with 0.4 mM Isopropyl β- d-1- thiogalactopyranoside (IPTG) and cells were grown for 2-3 hours at 37 °C with vigorous shaking. Cells were harvested by centrifugation at 4500 × g for 15 min and resuspended in Buffer A (50 mM NaH_2_PO_4_, 300 mM NaCl, 10 mM imidazole, 6 M urea, 1 mM dithiothreitol (DTT), pH 8.0). Cells were lysed on ice using an ultrasonic homogeniser and the soluble fraction was collected by centrifugation at 40,000 × g for 1 hour at 4 °C and loaded onto a 5 ml HisTrap HP column (Cytiva) equilibrated with Buffer A. The column was washed with 5 column volumes (CV) of Buffer A and eluted with a gradient with Buffer B (50 mM NaH_2_PO_4_, 300 mM NaCl, 500mM Imidazole, 6 M urea, 1 mM DTT, pH 8.0). Eluted samples were dialysed overnight against Buffer C (50 mM NaH_2_PO_4_, 150 mM NaCl, 2 M urea, 1 mM DTT, pH 8.0), followed by ULP1 protease (produced in-house) cleavage to remove the His6-SUMO tag. Following cleavage, Ascl1 constructs were loaded onto a 5 ml HisTrap HP column (Cytiva) to separate cleaved Ascl1 and 6xHis-SUMO tag. All protein preparations purity were checked by SDS-PAGE and pure fractions concentrated using Amicon Ultracentrifugal filters (Merck), reduced with DTT and purified by reversed-phase high-performance liquid chromatography (RP-HPLC) using a ZORBAX 300SB-C3 column (Agilent) with flow rate of 2.5 ml/min starting at 95% RP-HPLC solvent A 99.9% H_2_0, 0.08% trifluoroacetic acid (TFA)(Sigma) and 5% RP-HPLC solvent B (99.9% acetonitrile, 0.08% TFA) and going to 100% RP-HPLC solvent B over 95 min. Protein purity was analysed by SDS-PAGE, and samples were lyophilised and stored at −80 °C.

N-terminally hexahistidine-tagged E12 was expressed in LEMO21 cells with pET28b(+) vector cultured in LB-broth medium. Expression was induced at OD_600_ 0.6–0.8 with 1 mM IPTG and expressed for 4h at 37°C. Cells were harvested at 5000×g for 30 min at 4 °C. Cells thawed in 50 mM Tris, 150 mM NaCl, 0.2 mM EDTA, 0.5 mM DTT, pH 8 and lysed on ice using the ultrasonic homogeniser (Hielscher)and pelleted by centrifugation at 40,000×g for 1h at 4 °C. Lysis pellet resuspended in 6 M urea, 50 mM tris, 150 mM NaCl, 1 mM DTT, 5 mM imidazole, pH 8 and let incubate for 10-15 min at room temperature, centrifuged at 30000×g for 30 min at 4 °C and the supernatant was collected. The supernatant was purified by Ni^2+^ affinity chromatography. Eluted samples were pooled and dialysed concentrated by centrifugation in 50 mL Amicon spin filters with 30 kDa cut-off at 4000 × g and dialysed overnight against buffer A (50 mM Tris, 150 mM NaCl, 1 mM DTT, pH 8.0). The sample was further purified by reverse phase HPLC by a Reprosil Gold 300 C4 4.6×250 mm column and were analyzed by SDS-PAGE and finally by mass spectrometry. The purified samples were lyophilised and resuspended in 8M urea and used for interaction studies with labeled Ascl1 constructs.

### Protein labeling

The protocol for fluorescent labeling of proteins is based on published work from Zosel *et al*.^48^ The acceptor-donor dye pair used for protein labeling are Cy3B (Amersham) and CF660R (Biotium) modified with a reactive maleimide group. The reduced and lyophilized protein was dissolved in labeling buffer (0.1 M potassium phosphate, 6 M GdmCl, pH 7.2) and labeled at room temperature for 2 hours using Cy3B maleimide (donor) (0.7:1 dye to protein ratio). Unreacted dye was quenched using 100 mM DTT to reduce its affinity to the column and reduce oxidized cysteines. The reaction was incubated for 20 min at room temperature before purifying by RP-HPLC separate unlabeled and double donor-labeled proteins. For full length Ascl1, the ReprosilGold 200 C4 150×4.6mm was used, whereas the ReprosilGold 200 C18 250x4.6mm column was used for the shorter Ascl1 fragments. For the second labeling the purified protein fractions from the first step were pooled and lyophilised overnight, then resuspended in labeling buffer and labeled at RT for 2 hours using CF660R maleimide (1.3:1 dye to protein ratio). The reaction was quenched using DTT and RP-HPLC was then used to remove unreacted dye, and separate donor-donor doubly labeled, and acceptor-acceptor doubly labeled proteins. Donor-acceptor labeled proteins pooled and lyophilised, resuspended in 6 M GdmCl, frozen in liquid N2, and stored at −80 °C.

### DNA oligonucleotides

Single stranded (ss) DNA oligonucleotides for specific (E-box, 5’-ACTCCAGCTGTGGAT-3’) and nonspecific (lacking E-box, 5’-GATCGCCTCCCGCCT-3’) DNAs were purchased from Integrated DNA Technologies (IDT). To produce double stranded DNA (dsDNA) the individual oligonucleotides were dissolved in 10 mM tris, 50 mM NaCl, 1 mM EDTA pH 7.7 to a concentration which is twice the concentration of the desired dsDNA.

Equal volumes of the equimolar forward and reverse oligonucleotides were then mixed in a sterile Eppendorf tube. The annealing sample was incubated at 95°C for 5 min and then allowed to cool slowly down to room temperature. The DNA concentration was determined by measuring 260 nm absorbance.

### Single-molecule spectroscopy

All single molecule fluorescence experiments were conducted at 20 °C using a MicroTime 200 or Luminosa (PicoQuant, DE) connected to an Olympus IX73 inverted microscope. The donor and acceptor dyes were excited using a 520 or 530 nm diode laser (LDH-D-C-520, PicoQuant) and a 640 nm diode laser (LDH-D-C-640, PicoQuant) alternatively with a repetition rate of 40 MHz using pulsed interleaved excitation (PIE)^49^. The laser intensities were adjusted to 44 μW at 520 nm and 25 μW at 640 nm (PM100D, Thorlabs). Excitation and emission light was focused and collected using 60 × water objective (UPLSA-PO60XW, Olympus). Emitted fluorescence was focused through a 100 μm pinhole before being separated first by polarization and then by donor (582/64 BrightLine HC, Semrock) and acceptor (690/70 H Bandpass, AHF) emission wavelengths, into four detection channels. Detection of photons took place using single photon avalanche diodes (SPCM-AQRG-TR, Excelitas Technologies). The arrival time of detected photons was recorded with a MultiHarp 150 P time-correlated single photon counting (TCSPC) module (PicoQuant). All experiments were performed in μ-Slide sample chambers (Ibidi) at RT in TEK buffer (10 mM Tris, 0.1 mM EDTA, pH 7.4) with varying KCl concentrations. For photoprotection 143 mM 2-mercaptoethanol (Sigma) was added, along with 0.01% (v/v) Tween-20 (AppliChem) to reduce surface adhesion. In experiments using denaturants, the exact concentration of denaturant was determined from measurement of the solution refractive index.

### Analysis of transfer efficiency histograms

Data for transfer efficiency histograms were collected from 50 to 200 pM concentrations of double labeled freely diffusing Ascl1. Analysis was performed using the “Fretica” Mathematica scripting package (https://schuler.bioc.uzh.ch/programs/), created by Daniel Nettels and Ben Schuler. Fluorescence bursts were identified by the ΔT method, with interphoton durations under 100 µs and a minimum number of 30–40 photons per burst. Transfer efficiencies for each burst were computed as 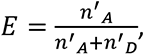 with n’A and n’D denoting the counts of acceptor and donor photons, respectively, adjusted for background fluorescence, direct acceptor excitation, cross-channel interference, and disparities in dye quantum yields and photon detection efficiencies^50^. Bursts affected by acceptor bleaching during transit through the confocal volume were excluded to avoid underestimating FRET. Bursts significantly exceeding the average photon counts, indicative of aggregates, were also removed. The stoichiometry ratio (*S*) for labeling was calculated using:

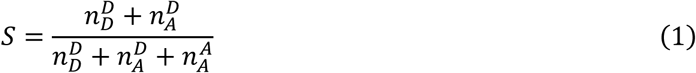

where 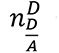 is the count of detected donor or acceptor photons post donor excitation and 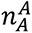 is the count of acceptor photons post acceptor excitation. To create the final transfer efficiency histograms, we selected bursts that have S=0.3-0.8; this selection helped isolate bursts from incorrectly labeled molecules, discarding those without an active acceptor. Despite filtering, a large donor only population can cause residual donor only bursts to remain in final histograms. To calculate average FRET efficiencies, we applied Gaussian or lognormal distribution fits to the histograms, identifying distinct populations within. The shot noise limited upper bound width of the distributions (**Supplementary Fig. S3)** were calculated based on Gopich et al.^51^. and depends on minimum number of photons in bursts chosen as the burst selection cut-off according to *σ*^2^_shot−noise_ ≤ 〈E〉(1 − 〈E〉)/N_T_ where NT is the minimum number of photons in a burst.

For comprehensive binding affinity analysis, multiple histograms were fitted globally, allowing certain parameters to be consistent across different measurements. Distance metrics from FRET efficiency data utilized the Förster resonance energy transfer equation:

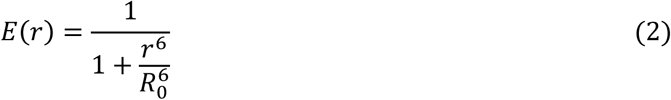

Where *R*_0_ is the Förster radius and *R*_0_ = 6.0 nm was calculated for the Cy3B/CF660R dye pair for a solution having refractive index near equal to 1.333. For denaturation and salt dependency experiments the *R*_0_ was corrected for changes in the refractive index of the solution measured at room temperature by a digital Abbe refractometer for each denaturant and salt concentrations according to the equation previously described^52^:

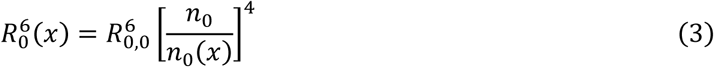

where *n*(x) is the refractive index of the sample at condition x.

For double labeled Ascl1 involving disordered regions we converted mean transfer efficiencies 〈*E*〉 to root-mean-square end-to end distances 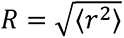 by numerically solving the following transcendental equation:

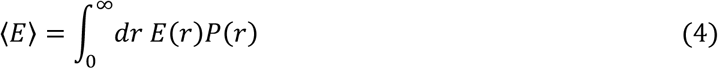

Here, *P*(*r*) denotes the distance probability density function of the SAW-*ν* model:

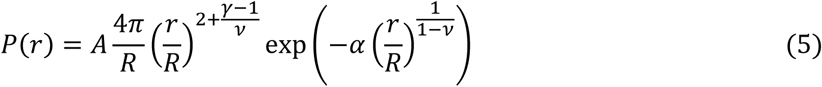

which is characterized by the critical exponents, *ν* and *γ* ≈ 1.1615. The constants *A* and *α* are determined by forcing *P*(*r*) to be normalized and to satisfy 〈*r*^2^〉 = *R*^2^. By adopting a scaling law *R* = *b N^v^*, we eliminate the *ν* dependency from *P*(*r*), substituting it with the expression 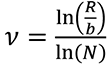 where *b* ≈ 0.55 corresponding an average value of approximately 0.55 nm for proteins and *N* indicates the number of monomer units separating the fluorescent dyes.

### Fluorescence correlation spectroscopy

All fluorescence correlation spectroscopy measurements were done with the same instrument and similar buffer and protein concentration as the smFRET measurement. We performed by correlating the intensity fluctuations in fluorescence in an smFRET experiment according to:

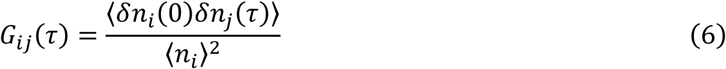

where where i,j=A, D and n_i_(0) and n_j_(τ) are fluorescence count rates for channels i and j at time 0 and after a lag time τ, respectively, and *δn_i_*_,*j*_ = *n_i_*_,*j*_ − 〈*n_i_*_,*j*_〉 are the corresponding deviations from the mean count rates. The donor-donor, acceptor-acceptor and cross-correlation curves were fitted with a three-dimensional free diffusion model with one diffusional component and another component for triplet blinking using the following equation:

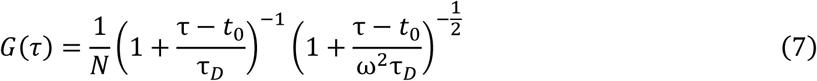

where N is the average number of molecules in the confocal volume, τ_D_ is the translational diffusion time, and ω = ω*_z_*/ω*_xy_* is the axial ratio of the confocal volume.

The nsFCS^22^ measurements were carried out with the 520 nm diode laser operating in continuous-wave mode at a laser power of 100 μW. The emitted photons were split by either a 50/50 beam splitter or polarizing beam splitter before being separated and filtered based on wavelength and detected at respective donor and acceptor detectors. The typical data acquisition times for nsFCS measurements were 16–20 h. All measurements were performed at protein concentrations of ∼1–2 nM, in TEK buffer similar to the smFRET measurements, 0.01% Tween-20, 143 mM β-mercaptoethanol. Donor and acceptor fluorescence photons from only the FRET subpopulation were used for correlations at 1 ns binning time. Photons were cross-correlated between detectors to avoid the effects of detector dead times and after-pulsing on the correlation functions. The auto-correlation curves of donor and acceptor channels and cross-correlation curves between donor and acceptor channels were calculated, fit, and analyzed as described previously^53^. Briefly, the correlation curves were fit over lag time interval from −1 μs to +1 μs using:

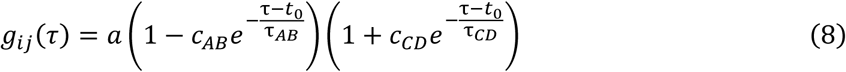

where *i* and *j* indicate donor (D) or acceptor (A) fluorescence emission; the amplitude a depends on the effective mean number of molecules in the confocal volume and on the background signal; *c*_AB_, *τ*_AB_, *c*_CD_, and *τ*_cd_, are the amplitudes and time constants of photon antibunching (AB), chain dynamics (CD), and triplet state of the dyes respectively. An additional term 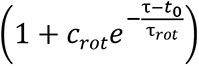 was also included for fitting the correlation curves for Ascl1 in presence of E12 to describe the fast decay observed on a short timescale (∼10 ns). The timescale, τ_rot_, can be associated with the restricted dye mobility in the complex because of the asymmetry between the photon correlations for positive and negative lag times when a polarizing beam splitter is used to separate the two major channels of detection was observed specially for the Ascl1-E12 complex (**Supplementary Fig. S5**)^54^, although an increase in static quenching of the dyes by aromatic residues of E12 in the complex can’t be ruled out^55,56^. The three correlation curves from each measurement were fit globally. τ_CD_ can be converted to the reconfiguration time of the chain, τ_r_ as described by Gopich et al.^23^, by assuming that the chain dynamics can be modelled as a diffusive process in the potential of mean force *F*(*r*) = −*k_B_T lnP*(*r*) derived from the sampled inter-dye distance distribution *P(r)* based on the SAW-ν model. The correlation functions are displayed with a normalisation to 1 at their respective values at 0.5 μs. The reported uncertainty of the reconfiguration time is a systematic error of the global fit.

### Fluorescence lifetime analysis

Fluorescence lifetimes were derived from the average detection times of the donor 〈*t_D_*〉 following its excitation pulse. These lifetimes were subsequently mapped against their corresponding transfer efficiencies on two-dimensional scatter plots. Here, 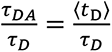 was determined for each fluorescence burst given an inherent donor lifetime *τ_D_*. In scenarios where donor-acceptor distance remains constant, the ratio 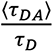 must match 1 − *E*. However, in systems exploring a wide range of distances, governed by a probability density function *P*(r) for the inter dye distance *r*, the mean fluorescence lifetime 〈*τ_DA_*〉 is influenced by the distribution of distances as according to:

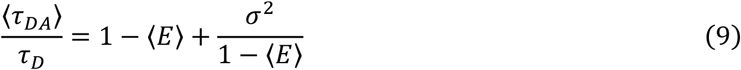

Here, the variance *σ*^2^ is given by:

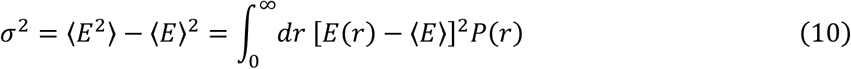

### Binding affinity measurements

Transfer efficiency histograms for doubly labeled Ascl1 were collected as the concentration of the unlabeled counterpart was incrementally increased until a stable transfer efficiency was observed. The data were segmented into two distinct subpopulations, representing bound and unbound states, using Gaussian peak fitting. The proportion of the bound fraction (*θ*) was then determined based on the relative area under these peaks. To derive the dissociation constant (*K*_D_), we applied a binding isotherm model to the data, fitting it with the following equation:

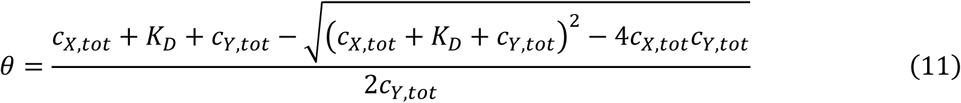

where *c_x_*_,*tot*_ and *c_Y_*_,*tot*_are the total concentration of binding partners, respectively, with one kept constant throughout the experiment.

### Analysis of chemical denaturation

For chemical denaturation, the dependence of the inter-dye end-to-end distance as a function of denaturant concentration was obtained by fitting to a weak binding model originally developed for change in FRET efficiency (*E*) with chemical denaturant^57^,

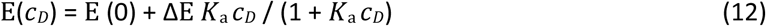

where *c_D_* is the denaturant concentration, *E*(0) is the FRET efficiency at zero denaturant, and *K*_a_ (association constant of the denaturant), ΔE, and are fit parameters. All the denaturant concentrations were obtained according to the refractive index of the solutions at each separate concentrations and to calculate the end-to-end inter dye distance the Förster radius (*R*_0_) were adjusted at each denaturant concentration due to the change in refractive index of the solutions. For the fitting of the denaturation of the heterodimeric complex between Ascl1-E12 the following two state protein unfolding model has been used as, 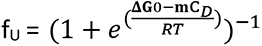 where *f*_u_ is the fraction of the dimer dissociated, *c_D_* is the denaturant concentration, Δ*G*_0_ is the folding free energy, *R* is the universal gas constant (0.001987 kcal/mol/K) and T set at 298K.

### System for studying protein abundance

The cell line used for the microscopy and flow cytometry experiments is the HEK293T TetBxb1BFPiCasp9 Clone 12 cell line^58^. The plasmid containing the NLS-GFP-fused Ascl1 variants also includes an internal ribosomal entry site (IRES), mCherry and a site for Bxb1 recombination into a landing pad in HEK293T cells. Its expression from the landing pad is controlled by a Tet-on promoter, that in non-recombined cells drives the expression of BFP, inducible Caspase 9 (iCasp9) and a blasticidin resistance gene separated with a parechovirus 2A-like translational stop-start sequence. When correct integration occurs, the GFP-fragments and mCherry are expressed from the same mRNA, allowing normalization of the GFP levels. The transfected Ascl1 constructs were made by Genscript.

### Protein abundance in HEK293T cells using flow cytometry

Cells were grown in Dulbeccós Modified Eaglés Medium (DMEM) (Sigma-Aldrich) with 10% (v/v) fetal bovine serum (FBS) (Sigma Aldrich), 0.24 mg/mL streptomycin sulphate (BioChemica), 0.29 mg/mL penicillin G potassium salt (BioChemica), 0.32 mg/mL L-glutamine (Sigma Aldrich) and 2 µg/mL doxycycline (Dox) (Sigma Aldrich). Cells were passaged upon reaching 70-80% confluency (passaging every two to three days). Cells were dislodged using trypsin (Gibco). The authenticity of the cell line was ensured by regular selection of recombinant cells with 10 nM of AP1903 (MedChemExpress) and based on expression of BFP in the non-recombinant cells. The cells tested negative for mycoplasma (Mycostrip, InvivoGen).

Transfections were performed in 12-well plates with 0.2 × 10^6^ cells. The Ascl1 constructs were mixed with the pCAG-NLS-Bxb1 (Addgene, Plasmid #51271) plasmid at a molar ratio of 17.5:1 along with OptiMEM (Thermo Fisher Scientific) (80 μL) and Fugene HD (Promega) (1.7 μL) and added to the cells. After 48 h Doxycyclin (2 μg/mL) was added in the cell culture, as well as 10 nM of AP1903 to ensure counterselection of non-recombinant cells. The cells were grown for 3 days before flow cytometry was performed. On day 3 after Dox addition, cells were treated with 15 μM of bortezomib (BZ) or the equivalent amount of DMSO for 16 h. Flow cytometry was performed on day 4 after Dox on a BD FACSCelesta flow cell cytometer. Cells were washed with PBS and dislodged with trypsin. Trypsin was deactivated by addition of DMEM full medium (with FBS) and the cells were centrifuged at 300 × g for 5 min. The cell pellet was resuspended in 500 μL of PBS with 2% FBS, which was the solution used for flow cytometry. For excitation of GFP, mCherry and BFP 488 nm, 561 nm and 405 nm lasers were used respectively. The filters used were 530/40, 610/20 and 450/40 respectively. All flow cytometry experiments were performed in duplicates. Gating strategy is included in the supplemental material (**Supplementary Fig. S12**).

### Fluorescence imaging

To track cellular localization and aggregation of Ascl1 we used the Zeiss AX10 microscope with a 10× objective lens. The preparation of the samples was identical to the one described for flow cytometry, without the steps for detaching the cells from the 12-well plates. The GFP and mCherry were excited by a 475 nm and a 590 nm led light from Colibri 7 respectively. All microscopy experiments were performed in duplicates.

## ACKNOWLEDGEMENTS

We would like to extend our deepest gratitude to Aida Curovic and Kristinn R. Óskarsson for technical assistance. We thank Birthe B. Kragelund for providing an expression plasmid for hexahistidine-tagged SUMO protein. We gratefully acknowledge funding from the European Research Council (ERC StG 101040601-PIONEER, to P.O.H.), the Icelandic Research Fund (grant no. 239556-051, to P.O.H.), and the Independent Research Fund Denmark, Technology and Production Sciences (grant no. 5284-00009B, to R.H.P.).

## DATA AVAILABILITY

Data supporting the findings of this paper are available from the corresponding authors upon reasonable request.

## CODE AVAILABILITY

Fretica, an add-on package for Mathematica v.15 (Wolfram Research), was used for the analysis of single-molecule data (available at https://schuler.bioc.uzh.ch/programs/).

## AUTHOR CONTRIBUTIONS

S.M., S.F.R., and P.O.H. conceived the study. S.M., S.F.R., Á.H.V.L., K.N., and A.G. purified all Ascl1 and E12 variants. S.M., S.F.R., Á.H.V.L., K.N., and A.G. performed all single-molecule experiments. S.M., S.F.R., Á.H.V.L., K.N., A.G., and P.O.H. analysed single-molecule data. V.V. conducted protein abundance and fluorescence imaging experiments. V.V. and R.H.P. analysed protein abundance data. K.S.Z. provided new reagents and protocols. S.M., V.V., and P.O.H. made the figures. S.M. and P.O.H. wrote the manuscript with input from all authors. P.O.H. was responsible for overall project management and supervised the research.

