## Supplementary Material for "PolyA/polyQ-mediated conformational rewiring regulates DNA engagement and drives aggregation in the neuronal transcription factor Ascl1"

```
Leopard         MESSAKMESGSAGQQPQPQP--------QPQQPFLPPAACFFATAAAAAAAAAAAAAQSAQQQQQQQQQQQQQQQQQQQQQQQQQQAPQLSPAA-DGQPSGGGHKSASKQVKRQRSSSPE    111
Bison           MESSAKMESGGAG------P--------QPQQPFLPPAACFFATAAA----------QSAQQP------------PPPPPPP-----PQLSPDGPTASPQGAVTSTAPKQVKRQRSSSPE    79
Bovine          MESSAKMESGGAG------P--------QPQQPFLPPAACFFATAAA----------QSAQQP------------PPPPP-----PPPQLSPA-ADGQPSGGGHKSAPKQVKRQRSSSPE    78
Sperm           MESSAKMESGGAG------P--------Q---PFLPPAACFFATAAAAAAAAAAAA-QSAQQP------------PPPPPPPPPPQAPQLSPA-ADGQPSGGGHKSAPKQVKRQRSSSPE    89
Galago          MESSAKMESGAVQQQPQPQPQPQPQPQSQPQPPFLQPTACFFATAAAAAAAAAAAAAA-------QSA--------------QHQQAGQLSPA-ADGQPSGGGHKSAPKQVKRQRSSSPE    98
Dog             -------------RSGRL----------SRPQPFLPPAACFFATAAAAA-------AQSAQQQPQPQP--------------QPQQAPQLSPA-ADGQPSGGGHKSASKQVKRQRSSSPE    75
Human           MESSAKMESGGAGQQPQP----------QPQQPFLPPAACFFATAAAAAAAAAAAAAQSAQQQQQQQQ--------------QQQQAPQLRPA-ADGQPSGGGHKSAPKQVKRQRSSSPE    95
Chimpanzee      MESSAKMESGGAGQQPQP----------QPQQPFLPPAACFFATAAAAAAAAAAAAAQSAQ----QQQ--------------QQQQAPQLRPA-ADGQPSGGGHKSAPKQVKRQRSS---    88
Mouse           MESSGKMESGA---GQQP----------QPPQPFLPPAACFFATAAAAAAAAAAA--AQSAQQQQPQA--------------PPQQAPQLSPV-ADSQPSGGGHKSAAKQVKRQRSSSPE    90
Rat             MESSGKMESGA---GQQP----------QPPQPFLPPAACFFATAAAAAAAAAAAAAQSAQQQQQQQA--------------PQQQAPQLSPV-ADGQPSGGGHKSAAKQVKRQRSSSPE    92
                                            .   *** *:*********                                         ** *    ..*.*.  .:* *********

Leopard         LMRCKRRLNFSGFGYSLPQQQPAAVARRNERERNRVKLVNLGFATLREHVPNGAANKKMSKVETLRSAVEYIRALQQLLDEHDAVSAAFQAGVLSPTISPNYSNDMNSMAGSPVSSYSSD    231
Bison           LMRCKRRLNFSGFGYSLPQQQPAAVARRNERERNRVKLVNLGFATLREHVPNGAANKKMSKVETLRSAVEYIRALQQLLDEHDAVSAAFQAGVLSPTISPNYSNDMNSMAGSPVSSYSSD    199
Bovine          LMRCKRRLNFSGFGYSLPQQQPAAVARRNERERNRVKLVNLGFATLREHVPNGAANKKMSKVETLRSAVEYIRALQQLLDEHDAVSAAFQAGVLSPTISPNYSNDMNSMAGSPVSSYSSD    198
Sperm           LMRCKRRLNFSGFGYSLPQQQPAAVARRNERERNRVKLVNLGFATLREHVPNGAANKKMSKVETLRSAVEYIRALQQLLDEHDAVSAAFQAGVLSPTISPNYSNDMNSMAGSPVSSYSSD    209
Galago          LMRCKRRLNFSGFGYSLPQQQPAAVARRNERERNRVKLVNLGFATLREHVPNGAANKKMSKVETLRSAVEYIRALQQLLDEHDAVSAAFQAGVLSPTISPNYSNDLNSMAGSPVSSYSSD    218
Dog             LMRCKRRLNFSGFGYSLPQQQPAAVARRNERERNRVKLVNLGFATLREHVPNGAANKKMSKVETLRSAVEYIRALQQLLDEHDAVSAAFQAGVLSPTISPNYSNDLNSMAGSPVSSYSSD    195
Human           LMRCKRRLNFSGFGYSLPQQQPAAVARRNERERNRVKLVNLGFATLREHVPNGAANKKMSKVETLRSAVEYIRALQQLLDEHDAVSAAFQAGVLSPTISPNYSNDLNSMAGSPVSSYSSD    215
Chimpanzee      ----------SGFGYSLPQQQPAAVARRNERERNRVKLVNLGFATLREHVPNGAANKKMSKVETLRSAVEYIRALQQLLDEHDAVSAAFQAGVLSPTISPNYSNDLNSMAGSPVSSYSSD    198
Mouse           LMRCKRRLNFSGFGYSLPQQQPAAVARRNERERNRVKLVNLGFATLREHVPNGAANKKMSKVETLRSAVEYIRALQQLLDEHDAVSAAFQAGVLSPTISPNYSNDLNSMAGSPVSSYSSD    210
Rat             LMRCKRRLNFSGFGYSLPQQQPAAVARRNERERNRVKLVNLGFATLREHVPNGAANKKMSKVETLRSAVEYIRALQQLLDEHDAVSAAFQAGVLSPTISPNYSNDLNSMAGSPVSSYSSD    212
                          ***********************************************************************************************:**************

Leopard         EGSYDPLSPEEQELLDFTNWF    252
Bison           EGSYDPLSPEEQELLDFTNWF    220
Bovine          EGSYDPLSPEEQELLDFTNWF    219
Sperm           EGSYDPLSPEEQELLDFTNWF    230
Galago          EGSYDPLSPEEQELLDFTNWF    239
Dog             EGSYDPLSPEEQELLDFTNWF    216
Human           EGSYDPLSPEEQELLDFTNWF    236
Chimpanzee      EGSYDPLSPEEQELLDFTNWF    219
Mouse           EGSYDPLSPEEQELLDFTNWF    231
Rat             EGSYDPLSPEEQELLDFTNWF    233
                *********************
```

**Supplementary Figure S1**. **Sequence alignment of Ascl1 across mammals.** Overall, Ascl1 is highly conserved but sequence alignment of the N-terminal half between closely related mammals reveals that the polyA/polyQ tract varies widely in length and composition. The alignment was generated by CLUSTAL Omega[1].

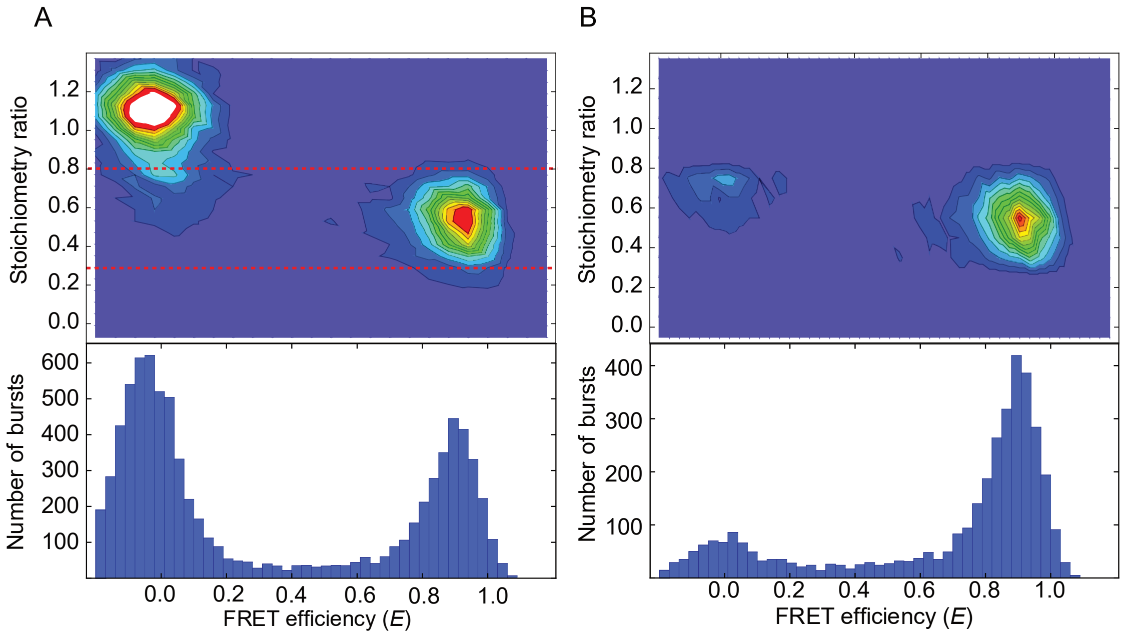


**Supplementary Figure S2**. **Stoichiometry measurements using pulsed-interleaved excitation.** Filtering for molecules that have stoichiometry close to 0.5 enables removing contributions from donor only-molecules from the FRET histogram. In cases where labelled samples had an excess of donor-only molecules, PIE filtering can be incomplete and leave a residual donor-only population. The stoichiometry graph and corresponding FRET efficiency histogram on the left is before applying filtering. After filtering (right), a residual donor-only population with FRET close to 0 persists.

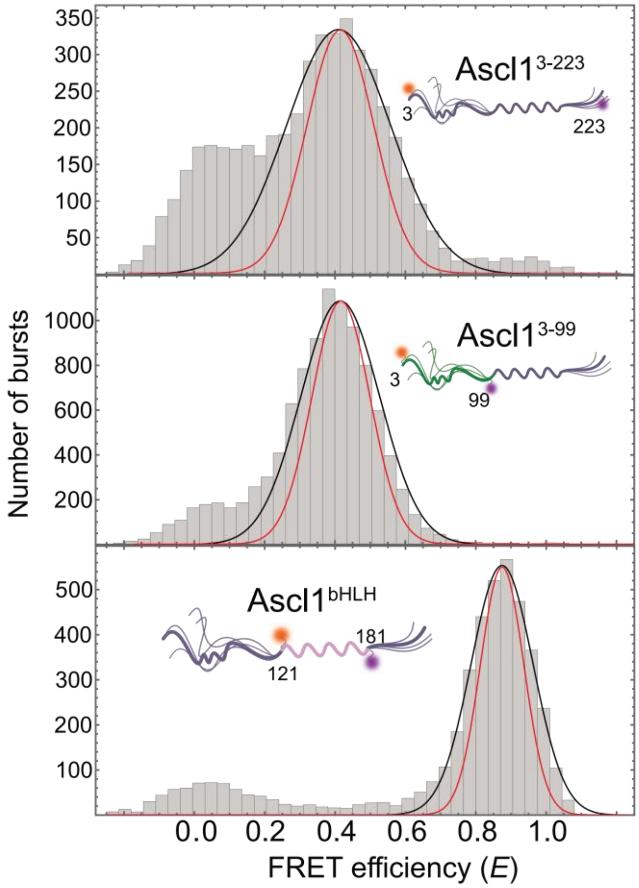


**Supplementary Figure S3. Peak broadening in single-molecule FRET efficiency histograms.** Photon distribution analysis was used to estimate shot-noise limited distribution widths (red fit lines) for Ascl1$^{3-223}$, Ascl1$^{3-99}$, and Ascl1$^{bHLH}$. Solid black lines are fits to a Gaussian distribution. Excess broadening beyond photon statistics was observed for all labelled variants and may be indicative of heterogeneous conformational sampling. See Methods for details.

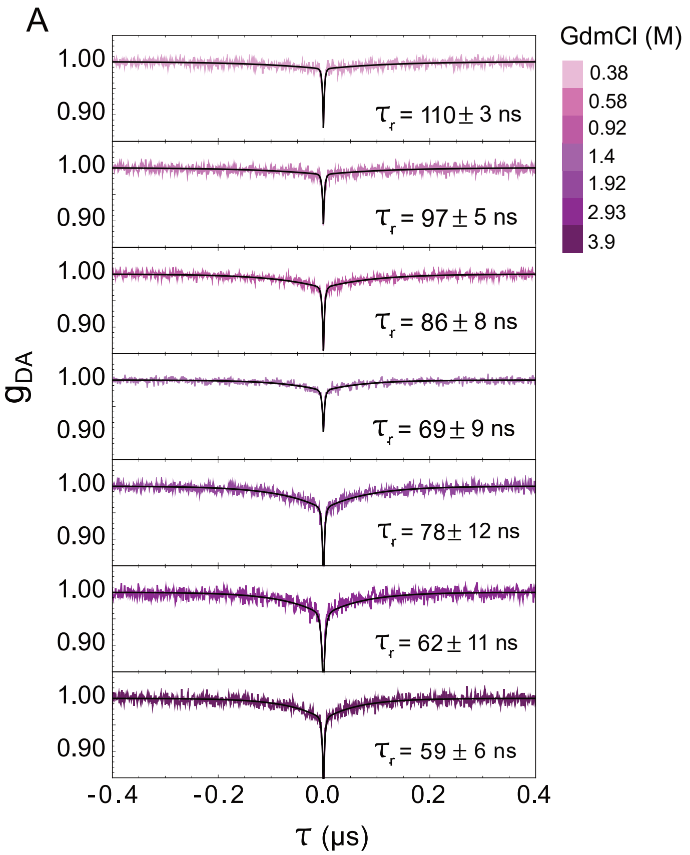


**Supplementary Figure S4. Nanosecond FCS analysis of Ascl1$^{bHLH}$ at different concentrations of GdmCl.** **A)** nsFCS correlation data for Ascl1$^{bHLH}$ at increasing concentrations of GdmCl. **B)** Reconfiguration time, $\tau_R$, for Ascl1$^{bHLH}$ as a function of GdmCl concentration.

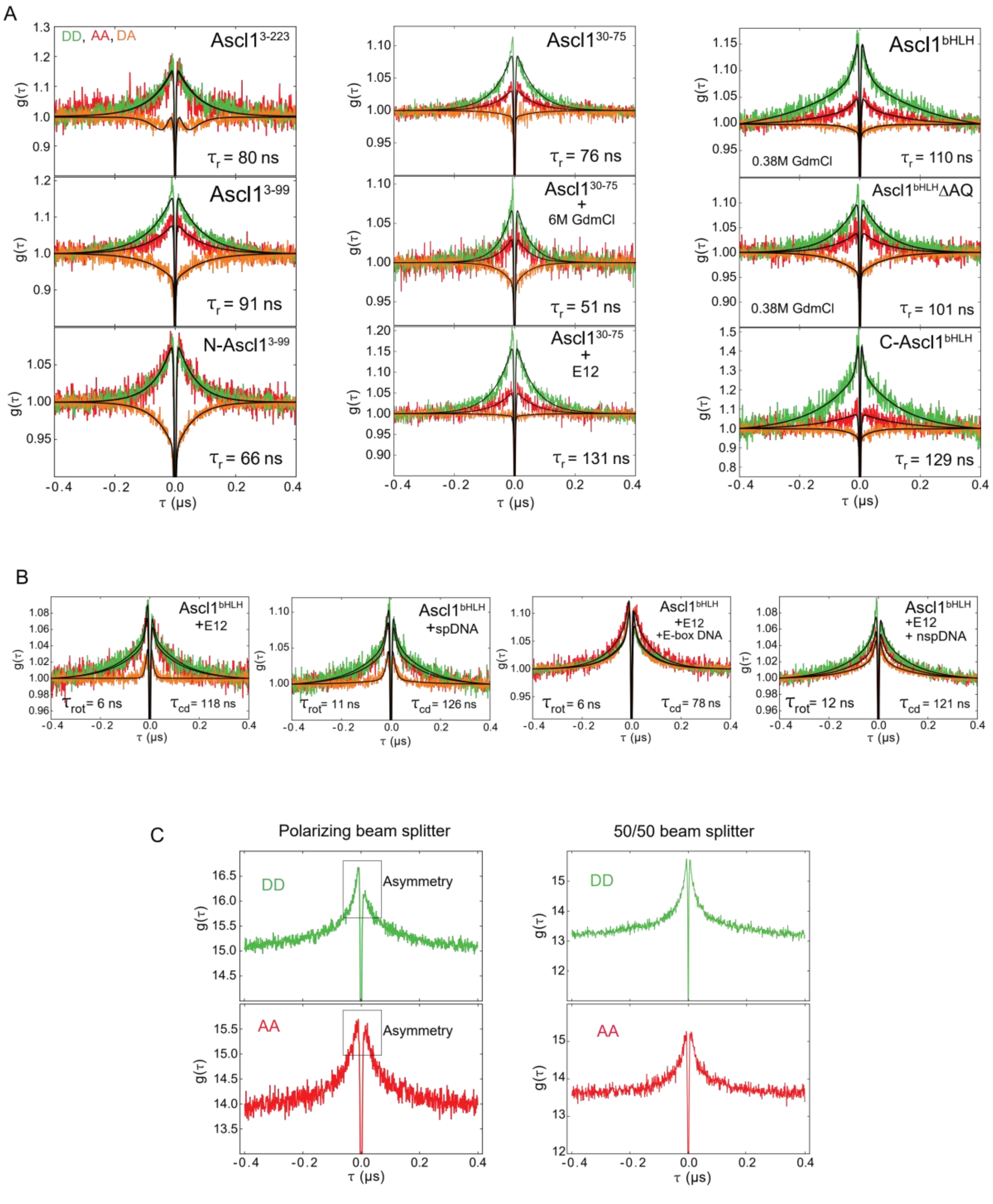


**Supplementary Figure S5. Full nsFCS correlation data analysis of Ascl1 constructs and their complexes at different conditions.** Donor-donor (green), acceptor-acceptor (red) and donor-acceptor (orange) correlation curves were acquired at 165 mM KCl concentration. The solid lines are global fits using Eq.8. **A)** Full nsFCS data for Ascl1 variants. **B)** nsFCS data for Ascl1$^{bHLH}$ in different molecular contexts. **C)** The effect of beam splitters on the asymmetry of the correlation curves.

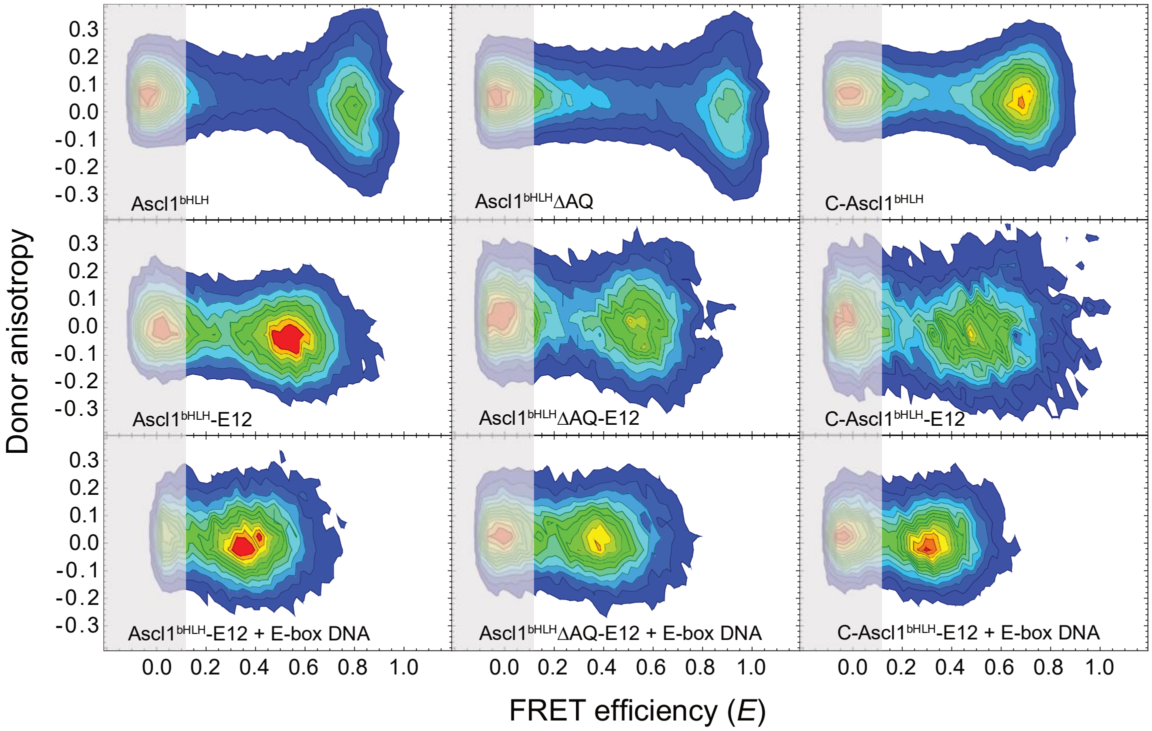


**Supplementary Figure S6. Steady state donor anisotropy plots of Ascl1 variants.** Steady-state donor anisotropy of the Ascl1$^{bHLH}$ variants in free, complex with E12, and heterodimeric complex with E-box DNA (indicated). The population near zero FRET efficiency is from the donor only population after PIE filtering. For all variants, the steady-state anisotropies are concentrated between -0.2 and 0.2, indicating rotational freedom of the donor dye[2,3].

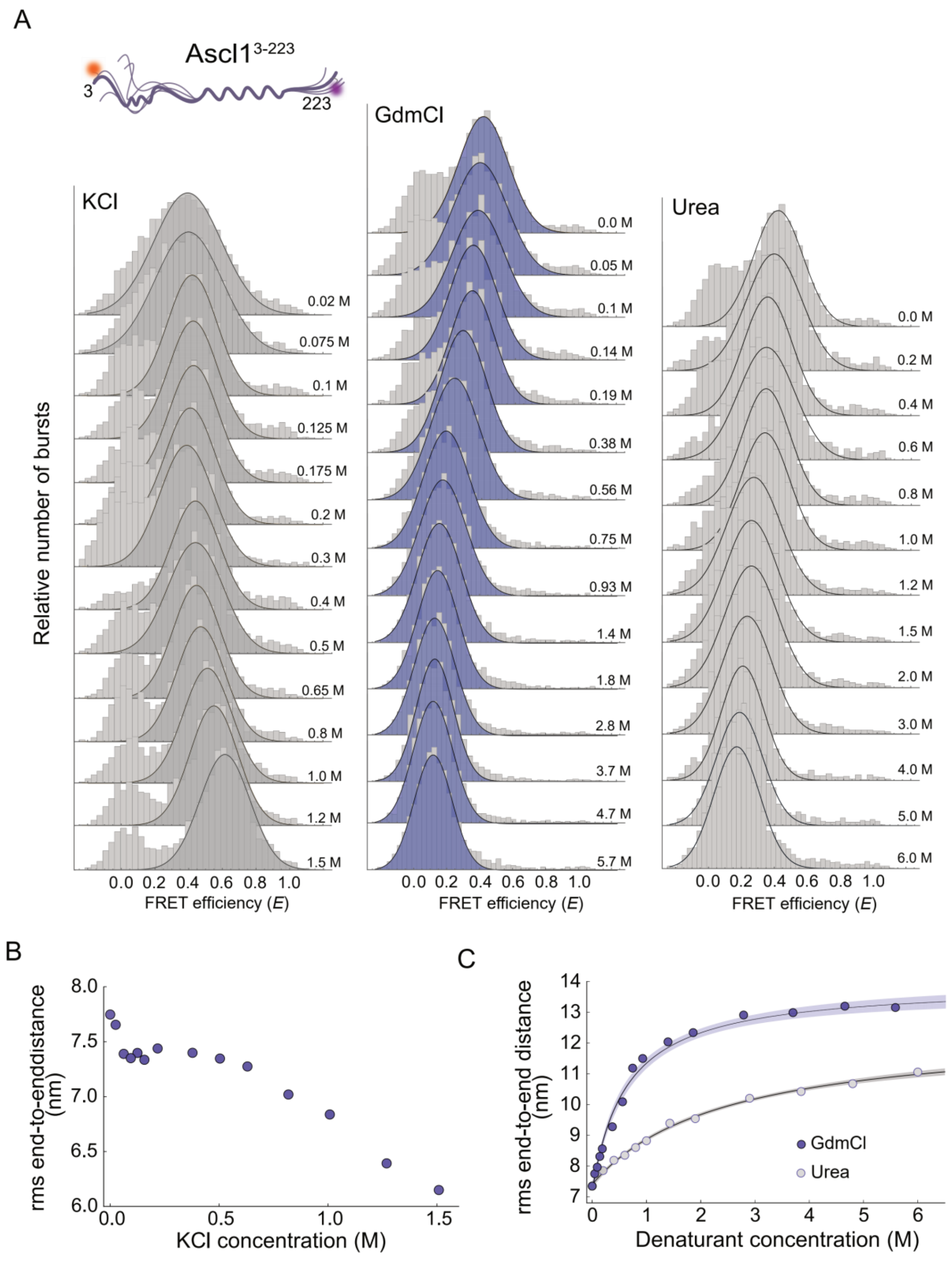
A
Ascl1[3-223]
3
223
KCl
GdmCl
Urea
Relative number of bursts
FRET efficiency (E)
B
rms end-to-enddistance (nm)
KCl concentration (M)
C
rms end-to-end distance (nm)
Denaturant concentration (M)
GdmCl
Urea

**Supplementary Figure S7. Ascl1$^{3\text{-}223}$ dimensions as a function of salt or denaturant concentration. A)** Single-molecule FRET efficiency histograms as a function of KCl concentration (left), GdmCl concentration (middle), and urea concentration (right). The solid lines are fits to a single Gaussian distribution. Each histogram consists of FRET efficiencies measured on >10000 molecules. **B)** RMS end-to-end distance as a function of KCl concentration. **C)** RMS end-to-end distance as a function of denaturant concentration (dark blue: GdmCl; light blue: urea). The shaded areas represent 95% confidence intervals. A complex pattern of compaction/expansion is observed with increasing KCl, whereas increasing denaturants leads to a gradual expansion of the polypeptide chain.

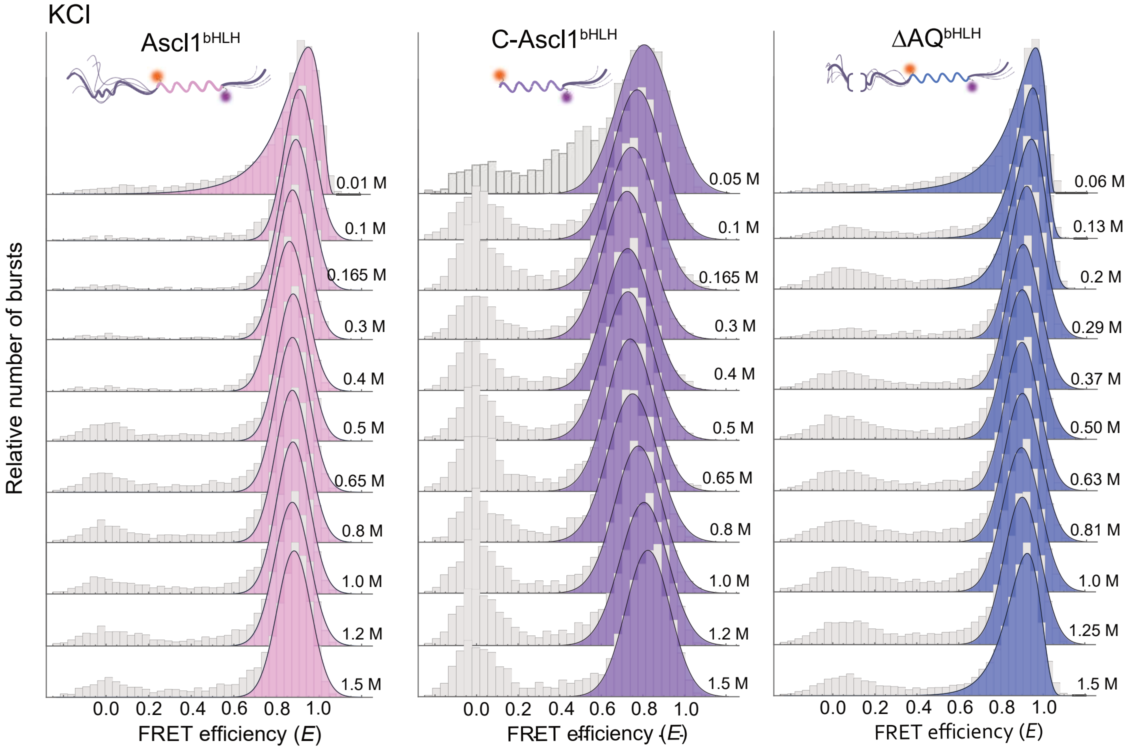


**Supplementary Figure S8**. **Single-molecule FRET efficiency histograms of Ascl1 variants as a function of KCl concentration. Left:** Ascl1$^{bHLH}$, **middle:** C-Ascl1$^{bHLH}$, **right:** $\Delta$AQ$^{bHLH}$. The solid lines are fits to a single Gaussian distribution. Each histogram consists of FRET efficiencies measured on >10000 molecules.

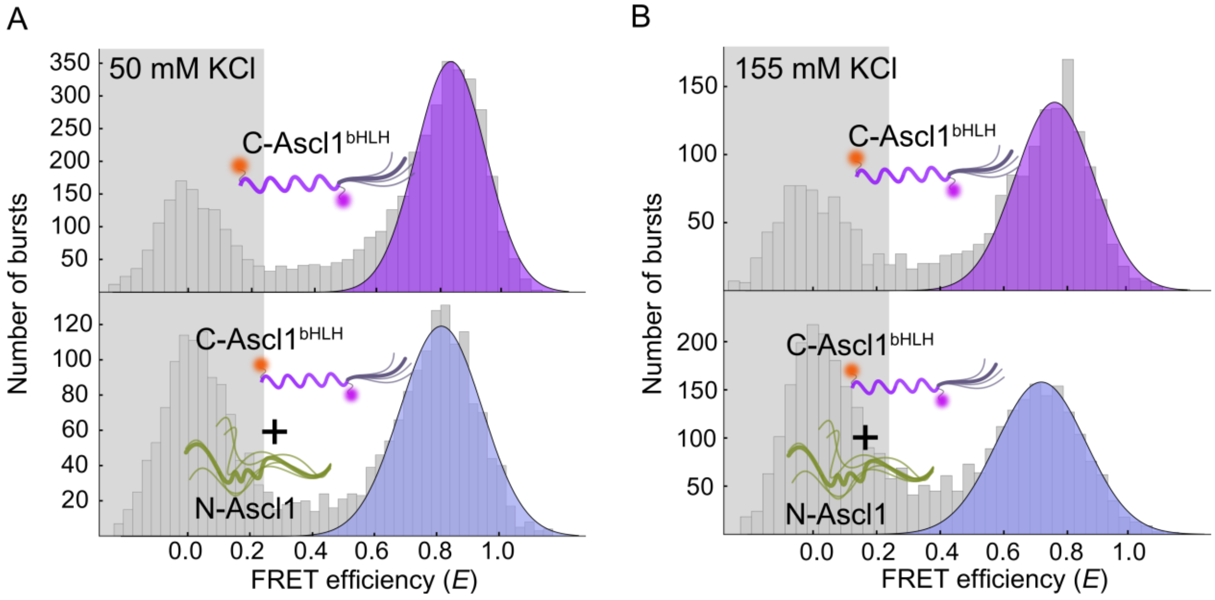


**Supplementary Figure S9. Interdomain interactions in Ascl1 are weak.** Single-molecule FRET efficiency histograms of fluorescently labelled C-Ascl1$^{bHLH}$ at **A)** 50 mM KCl, and **B)** 155 mM KCl, in the absence (upper) and presence (lower) of 2 µM of N-Ascl1.

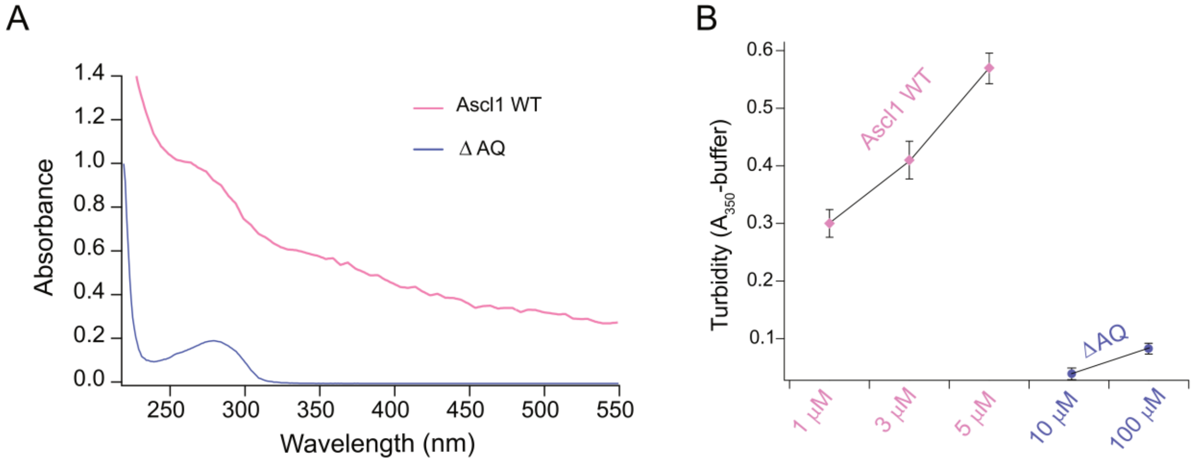


**Supplementary Figure S10. Turbidity measurements at 350 nm to monitor aggregation. A)** Absorbance spectra are consistent with aggregates at 5 µM concentration of full-length Ascl1 (grey lines) but a fully soluble ΔAQ at 10 µM. B) Turbidity (A350 nm - buffer) of Ascl1 full length samples at 1-5 µM and Ascl1ΔAQ samples at 10-100 µM total protein concentration in TEK buffer (10 mM Tris, 155mM KCl, 0.1 mM EDTA, 1mM DTT, pH 7.4) after 1 h incubation on ice. For Ascl1 full-length construct the purified protein was diluted from a 4 M GdmCl storage buffer stock into the experimental buffer to the final protein concentration.

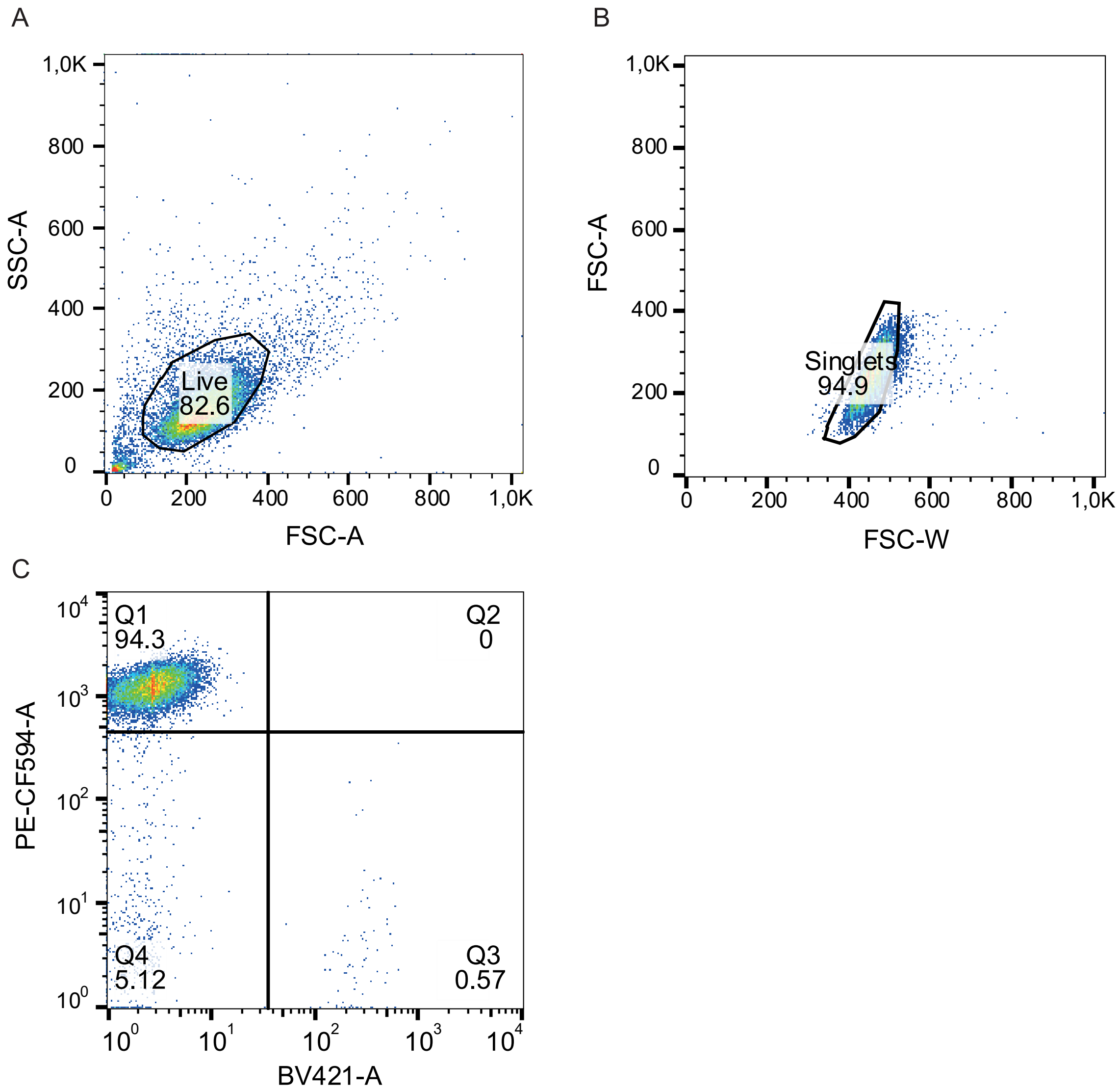


**Supplementary Figure S11. Flow cytometry gating strategy for recorded events. A)** Gating of live cell population based on side and forward scatter (SSC, FSC, respectively). **B)** Gating of singlets based on the forward scatter pulse area (FSC-A) and width (FSC-W). **C)** Gating of the BFP (BV421-A) negative and mCherry (PE-CF594-A) positive cells (recombinant cells).

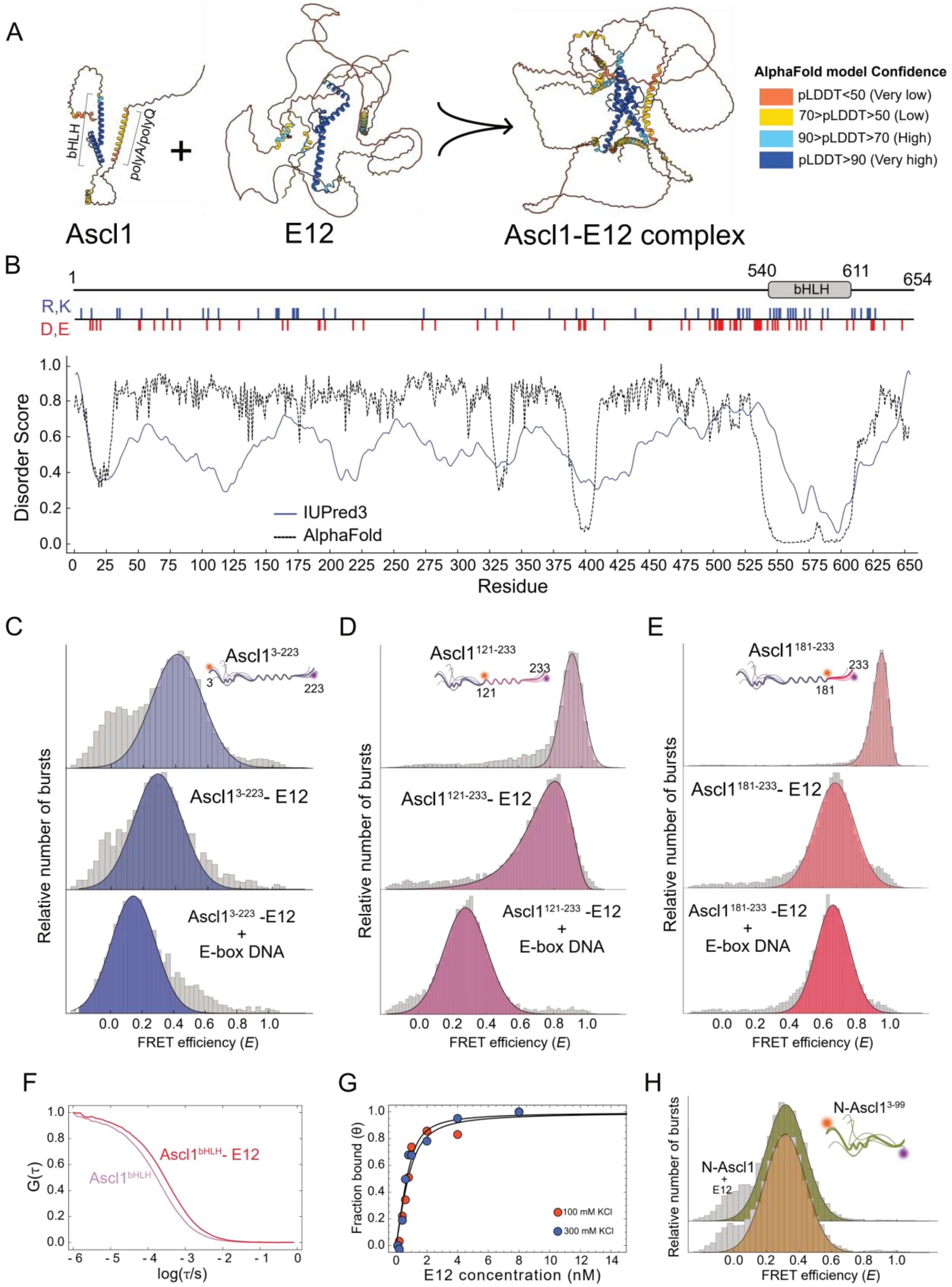
A
bHLH
polyA/polyQ
Ascl1
E12
Ascl1-E12 complex
AlphaFold model Confidence
pLDDT<50 (Very low)
70>pLDDT>50 (Low)
90>pLDDT>70 (High)
pLDDT>90 (Very high)
B
1
540
611
654
bHLH
R,K
D,E
Disorder Score
IUPred3
AlphaFold
Residue
C
D
E
Ascl1 3-223
Ascl1 3-223 - E12
Ascl1 3-223 -E12 + E-box DNA
Ascl1 121-233
Ascl1 121-233 - E12
Ascl1 121-233 -E12 + E-box DNA
Ascl1 181-233
Ascl1 181-233 - E12
Ascl1 181-233 -E12 + E-box DNA
Relative number of bursts
FRET efficiency (E)
F
G(τ)
log(τ/s)
Ascl1 bHLH - E12
Ascl1 bHLH
G
Fraction bound (θ)
E12 concentration (nM)
100 mM KCl
300 mM KCl
H
N-Ascl1 3-99
N-Ascl1 + E12
Relative number of bursts
FRET efficiency (E)

**Supplementary Figure S12. Properties of the Ascl1/E12 heterodimer. A)** AF3 predictions of Ascl1 and E12 monomers and their heterodimeric complex[4]. The colors represent pLDDT confidence intervals (Blue: high confidence, red: low confidence). **B)** Disorder prediction plot for E12 and pattern of charged residues. The bHLH domain is indicated. **C)** Single-molecule FRET efficiency histograms of $Ascl1^{3\text{-}223}$ free, in complex with E12, and the trimeric complex with DNA. **D)** Single-molecule FRET efficiency histograms of $Ascl1^{121\text{-}233}$ free, in complex with E12, and the trimeric complex with DNA. **E)** Single-molecule FRET efficiency histograms of $Ascl1^{181\text{-}233}$ free, in complex with E12, and the trimeric complex with DNA. **F)** Donor-acceptor cross-correlation of fluorescently labelled $Ascl1^{bHLH}$, labelled in positions 121 and 181. Free Ascl1 (pink) has a relatively short diffusion time through the confocal volume. In the presence of 20 nM of the E12 (unlabelled) the diffusion time is increased (red) due to the increased size of the molecular complex. **G)** $Ascl1^{bHLH}$ binding isotherms for complex formation with E12, at 100 and 300 mM KCl. The solid lines are fits to a 1:1 binding model. **H)** Single-molecule FRET efficiency histograms of N-$Ascl1^{3\text{-}99}$ in absence and presence of 8 nM E12. The lack of change in mean FRET efficiency supports that the N-IDR does not interact directly with E12.

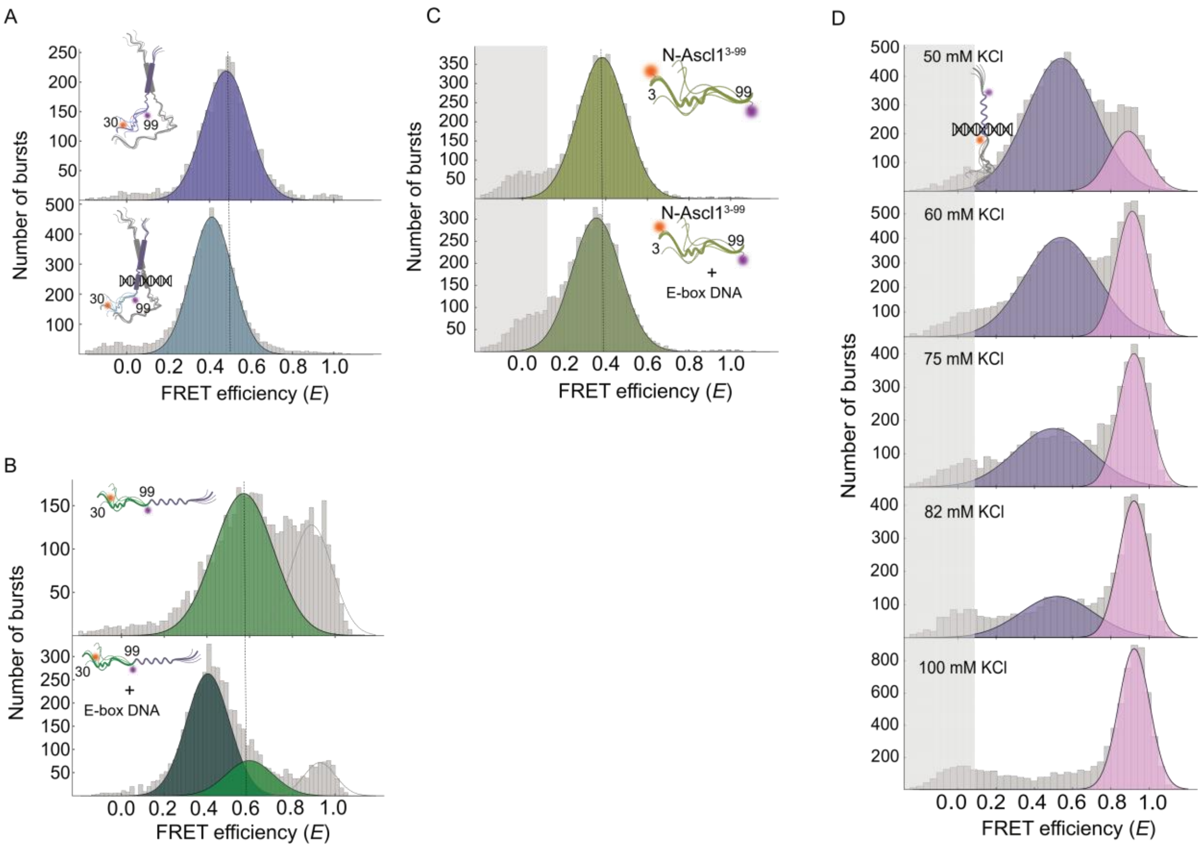


**Supplementary Figure S13**. **DNA binding characteristics of Ascl1. A)** Ascl1$^{30\text{-}99}$ in complex with E12 and after adding 1 μM E-box DNA. **B)** Ascl1$^{30\text{-}99}$ monomer E-box DNA interaction at 50mM KCl. **C)** N-Ascl1$^{3\text{-}99}$ with 20 μM E-box DNA at 50mM KCl. In the full-length molecule (panel B), the N-IDR expands upon DNA binding, but such an expansion is not observed with the isolated N-IDR. **D)** Single-molecule FRET efficiency histograms illustrating E-box DNA binding of Ascl1$^{bHLH}$ monomer at different salt concentrations, illustrating that monomer DNA binding is negligible at KCl concentrations above 100 mM.

**Supplementary Table 1**. Amino acid sequences of main protein constructs used in this work. Cysteine substitutions for dye labelling are indicated in bold red. For labelling positions involving residue 233, Tryptophan 235 (in bold) was mutated to phenylalanine to avoid complications from dye quenching.

| | |
|---|---|
| Ascl1 (WT) | MESSAKMESGGAGQQPQPQPQQPFLPPAACFFATAAAAAAAAAAAAAQSAQQQQQQQQQQQQAPQLRPAADGQP<br>SGGGHKSAPKQVKRQRSSSPELMRCKRRLNFSGFGYSLPQQQPAAVARRNERERNRVKLVNLGFATLREHVPNGAAN<br>KKMSKVETLRSAVEYIRALQQLLDEHDAVSAAFQAGVLSPTISPNYSNDLNSMAGSPVSSYSSDEGSYDPLSPEEQ<br>ELLDFTNWF |
| E12 | MNQPQRMAPVGTDKELSDLLDFSMMFPLPVTNGKGRPASLAGAQFGGSGLEDRPSSGSWGSGDQSSSSFDPSRTFSE<br>GTHFTESHSSLSSSTFLGPGLGGKSGERGAYASFGRDAGVGGLTQAGFLSGELALNSPGPLSPSGMKGTSQYYPSYSGS<br>SRRRAADGSLDTQPKKVRKVPPGLPSSVYPPSSGEDYGRDATAYPSAKTPSSTYPAPFYVADGSLHPSAELWSPPGQAG<br>FGPMLGGGSSPLPLPPGSGPVGSSGSSSTFGGLHQHERMGYQLHGAEVNGGLPSASSFSSAPGATYGGVSSHTPPVS<br>GADSLLGSRGTTAGSSGDALGKALASIYSPDHSSNNFSSSPSTPVGSPQGLAGTSQWPRAGAPGALSPSYDGGLHGLQ<br>SKIEDHLDEAIHVLRSHAVGTAGDMHTLLPGHGALASGFTGPMSLGGRHAGLVGGSHPEDGLAGSTSLMHNHAALPS<br>QPGTLPDLSRPPDSYSGLGRAGATAAASEIKREEKEDEENTSAADHSEEEKKELKAPRARTSPDEDEDDLLPPEQKAEREK<br>ERRVANNARERLRVRDINEAFKELGRMCQLHLNSEKPQTKLLILHQAVSVILNLEQQVRERNLNPKAACLKRREEEKVSG<br>VVGDPQMVLSAPHPGLSEAHNPAGHM |
| Ascl1$^{3-223}$<br>Ascl1$^{3-99}$<br>Ascl1$^{121-233}$<br>Ascl1$^{181-233}$<br>Ascl1$^{121-181}$<br>Ascl1$^{30-75}$<br>Ascl1$^{30-99}$ | 3 30<br>ME**C**SAKMESGGAGQQPQPQPQQPFLPPAA**C**FFATAAAAAAAAAAAAAQSAQQQQQQQQQQQQAPQLRPAADGQP<br>75 99 121<br>**C**GGGHKSAPKQVKRQRSSSPELMR**C**KRRLNFSGFGYSLPQQQPA**C**VARRNERERNRVKLVNLGFATLREHVPNGAAN<br>181 223<br>KKMSKVETLRSAVEYIRALQQLLDEHD**C**VSAAFQAGVLSPTISPNYSNDLNSMAGSPVSSYSSDEGSYDPL**C**PEEQELLD<br>233<br>F**C**N**W**F |
| N-Ascl1 | ME**C**SAKMESGGAGQQPQPQPQQPFLPPAACFFATAAAAAAAAAAAAAQSAQQQQQQQQQQQQAPQLRPAADGQP<br>SGGGHKSAPKQVKRQRSSSPELMR**C**KRRLNFSGFGYSLPQQ |
| C-Ascl1 | MQQP**C**AVARRNERERNRVKLVNLGFATLREHVPNGAANKKMSKVETLRSAVEYIRALQQLLDEHD**C**VSAAFQAGVLSP<br>TISPNYSNDLNSMAGSPVSSYSSDEGSYDPLSPEEQELLDFTNWF |
| $\Delta AQ^{bHLH}$ | MESSAKMESGGAGQQPQPQPQQPFLPPAACFFATAPQLRPAADGQPSGGGHKSAPKQVKRQRSSSPELMRCKRRLN<br>FSGFGYSLPQQQPA**C**VARRNERERNRVKLVNLGFATLREHVPNGAANKKMSKVETLRSAVEYIRALQQLLDEHD**C**VSAA<br>FQAGVLSPTISPNYSNDLNSMAGSPVSSYSSDEGSYDPLSPEEQELLDFTNWF |
| Ascl1$^{polyA/polyQ}$ | MESSAKMESGGAGQQPQPQPQQPFLPPAA**C**FFATAAAAAAAAAAAAAQSAQQQQQQQQQQQQAPQLRPAADGQP<br>**C**GGGHKSAPKQVKRQRSSSPELMRCKRRLNFSGFGYSLPQQQPAAVARRNERERNRVKLVNLGFATLREHVPNGAAN<br>KKMSKVETLRSAVEYIRALQQLLDEHDAVSAAFQAGVLSPTISPNYSNDLNSMAGSPVSSYSSDEGSYDP LSP EEQ<br>ELLDFTNWF |

**Supplementary Table 2**. Measured FRET efficiencies and related parameters for all protein variants.

| Ascl1 variant | Mean FRET efficiency (E) | Dye position | Sequence separation (#a.a.) | Scaling exponent (ν) | $\tau_{CD}$ / $\tau_{R}$ (ns) | D/A lifetime (ns) | FRET lifetime |
|---|---|---|---|---|---|---|---|
| $Ascl1^{3-223}$ | 0.41 | 3/223 | 220 | 0.47 | 72.8/84.6 | 2.72/3.03 | 0.86 |
| $Ascl1^{3-99}$ | 0.39 | 3/99 | 96 | 0.51 | 82.9/94.2 | 2.75/3.05 | 0.79 |
| $Ascl1^{polyA/polyQ}$ | 0.65 | 30/75 | 45 | 0.58 | 60.6/65.9 | 2.76/3.07 | 0.72 |
| $Ascl1^{bHLH}$ | 0.88 | 121/181 | 60 | 0.44 | 99.7/110.1 | 2.75/3.08 | 0.66 |
| $Ascl1^{181-233}$ | 0.95 | 181/233 | 52 | 0.42 | - | 2.89/ | 0.47 |
| $Ascl1^{121-233}$ | 0.91 | 121/233 | 112 | 0.39 | 81.7/97.3 | 2.89/3.11 | 0.64 |
| N-$Ascl1^{3-99}$ | 0.32 | 3/99 | 96 | 0.56 | 60.2/68 | 2.62/3.07 | 0.87 |
| C-$Ascl1^{bHLH}$ | 0.74 | 121/181 | 60 | 0.52 | 117.3/132.5 | 2.72/3.09 | 0.73 |
| $\Delta AQ^{bHLH}$ | 0.92 | 121/181 | 60 | 0.43 | 88.3/98.1 | 2.68/3.11 | 0.60 |

**Supplementary Table 3**. Measured FRET efficiencies for all Ascl1 variants in complex with E12, E-box DNA, and non-specific DNA.

| **Ascl1 variant** | | **Mean FRET efficiency (E)** | | |
|---|---|---|---|---|
| | Dye placement | E12 | E-box DNA | Non-specific DNA |
| $Ascl1^{FL}$ | 3/223 | 0.29 | 0.18 | - |
| $Ascl1^{N\text{-}IDR}$ | 3/99 | 0.32 | 0.32 | - |
| $Ascl1^{polyA/polyQ}$ | 30/75 | 0.59 | 0.52 | - |
| $Ascl1^{181\text{-}233}$ | 181/233 | 0.67 | 0.65 | - |
| $Ascl1^{121\text{-}233}$ | 121/233 | 0.81 | 0.30 | - |
| $Ascl1^{bHLH}$ | 121/181 | 0.48 | 0.39 | 0.40 |

**Supplementary Table 4.** E12 and DNA binding affinities for Ascl1 constructs in their monomeric and heterodimeric forms. Errors represent standard errors from fits to the corresponding binding isotherms.

| | E12 | DNA | | | |
|---|---|---|---|---|---|
| | | Ascl1 Monomer (50 mM KCl) | | Ascl1/E12Heterodimer (165 mM KCl) | |
| **Protein construct** | $K_d$ (nM) | $K_d$ (nM) $DNA^{E\text{-}box}$ | $K_d$ (nM) $DNA^{nonspecific}$ | $K_d$ (nM) $DNA^{E\text{-}box}$ | $K_d$ (nM) $DNA^{nonspecific}$ |
| $Ascl1^{bHLH}$ | 0.33±0.06 | 77±10 | 50±3 | 0.88±0.3 | 86 ±7 |
| $C\text{-}Ascl1^{bHLH}$ | 0.43±0.03 | 2564±440 | 1058±119 | 1.5±0.2 | ND |
| $\Delta AQ^{bHLH}$ | 0.22±0.05 | 1851±207 | 1408±261 | 1.2±0.2 | 600 ±80 |
